# In Vivo Massively Parallel Reporter Assay Reveals Sequence Determinants of mRNA Localization in Astrocytes

**DOI:** 10.64898/2026.04.27.721172

**Authors:** S.K. Koester, K. Sakers, G. M. Rurak, S.P. Plassmeyer, K. McFarland White, S. Alves Ferreira Dias, P. Baird, D.J. Kornbluth, J.D. Dougherty

**Affiliations:** Department of Genetics, Washington University School of Medicine, Saint Louis, MO, USA; Department of Psychiatry, Washington University School of Medicine, Saint Louis, MO, USA; Department of Neuroscience, Washington University School of Medicine, Saint Louis, MO, USA; Intellectual and Developmental Disabilities Research Center, Washington University, St. Louis, MO 63130, USA

## Abstract

RNA localization and local translation are essential mechanisms to fine-tune spatiotemporal gene expression in the nervous system. However, efficiently assessing the thousands of possible sequence determinants of RNA localization is a challenge, particularly for cell types that only reach morphological maturity *in vivo*. Here, we developed an *in vivo* Massively Parallel Reporter Assay (MPRA), termed Synaptoneurosomal (SN)-MPRA to enable identification of sequence determinants of mRNA localization and local translation, and applied this to astrocytes. We evaluated multiple models of RNA localization for two locally translated astrocyte mRNAs, *Glt1* and *Sparc*, including increased transcript abundance, “zipcode” elements, and RNA secondary structure. Our results establish a high-throughput *in vivo* framework for identifying *cis*-regulatory sequences driving RNA localization and local translation, and suggest astrocytes use diverse mechanisms to regulate subcellular gene expression. More broadly, SN-MPRA offers a versatile platform to study RNA localization *in vivo*, where biological context and intercellular interactions are preserved.

## Introduction

To perform specific cellular functions, polarized or functionally compartmentalized cells may preferentially localize a subset of mRNAs for spatiotemporally regulated protein synthesis. This conserved mechanism, local translation, has been well-characterized in numerous organisms and polarized cell types, including yeast^1^, root hairs^2^, oocytes and developing embryos^3–5^, neurons^6–10^ and oligodendrocytes^11,12^. More recent work has found that other glial cells of the central nervous system (CNS), such as radial glia^13^, astrocytes^14–16^, and microglia^17^, also exhibit regulated local translation.

Astrocytes are morphologically complex glial cells in the CNS with numerous essential roles. Peripheral astrocyte processes (PAPs) tightly ensheathe synapses to regulate a host of synaptic functions – including synaptic formation, maturation, elimination, and transmission – forming what is known as the “tri-partite synapse”^18,19^. Remarkably, in mice a single cortical astrocyte can contact over 100,000 synapses through its PAPs, with this number growing to upwards of 2,000,000 in humans ^20,21^. It is therefore hypothesized that local translation facilitates PAPs performance of these functions, enabling individual PAPs to respond more independently to local extracellular cues. In support of this, several studies identified specific mRNAs that are relatively enriched in the PAPs compared to the soma, with similar findings in radial glia processes^13–16^. Examples of locally translated mRNAs include *Sparc*, which encodes an astrocyte-secreted factor involved in synaptic refinement^22^, and *Glt1* (*Slc1a2*), which encodes a glutamate transporter necessary for uptake of excess glutamate from synaptic clefts, preventing excitotoxicity^23^. Further, PAPs produce local translational responses to neuronal activity and undergo translational changes during fear conditioning that alter protein composition^16,24^. This suggests a potential role of astrocytic local translation in learning and memory, as well as in synaptic and circuit function in general^16^. However, the rules by which specific sequences become translationally enriched at these processes are unclear. Understanding what mediates specific RNAs to localize to PAPs and regulates their translation is crucial to understanding how PAPs perform their essential functions.

Localization of mRNAs to sub-cellular compartments in other cell types has been shown to be often mediated by the 3’ UTR through several mechanisms^25^. The classical model for regulation of mRNA delivery is via interactions between specific *cis-*regulatory sequences, or ‘zip-codes,’ with *trans-*regulatory factors, such as RNA binding proteins (RBPs) and microRNAs (miRNAs)^26^. Early evidence for such models includes studies characterizing the β-actin zipcode^27^, which directs localization of β-actin mRNA to the leading edge of fibroblasts through ZBP1 binding^28^, and the 3’ UTR of CamKIIa, which mediates its mRNA localization to dendrites^29^. Other mechanisms involve combinatorial logic, such as the necessity of both the RNA transport signal (RTS) and RNA localization region (RLR) in the 3’ UTR of myelin basic protein (MBP), which direct and transport its mRNA to the myelin compartment in oligodendrocytes^11^. Outside of sequence motifs, interactions between secondary structures, such as G-quadruplexes, and RBPs, such as FMRP, have been shown to mediate localization in neurons ^30^. Likewise, alternate mechanisms simply regulating RNA stability have been shown to strongly influence RNA localization in neurons, whether mediated by RBPs or other features of the transcript^31^.

In astrocytes, much less is understood about the specific features of mRNAs driving their localization to the PAPs or the mechanisms involved. While PAP-enriched transcripts have higher expression than PAP-depleted transcripts^15^, astrocytic processes constitute the majority of the cell’s volume^32^, which may explain the corresponding increase in abundance of PAP-enriched transcripts. Additionally, locally translated transcripts also contain longer 3’ UTRs with more stable secondary structures^15^. Further, motif enrichment analyses on the 3’ UTRs of locally translated genes showed an enrichment of Quaking Response Elements (QRE). However, deletion of the QREs from the *Sparc* 3’ UTR did not significantly impact peripheral RNA localization, but rather impacted nuclear export and local protein levels ^15^. Further, UTRs can be long (e.g., >9 Kb GLT1a 3’ UTR), thus harboring thousands of possible regulatory sequences to be evaluated. Therefore, a high-throughput method to systematically define RNA sequence motifs necessary and sufficient to drive mRNA localization in astrocytes is needed.

In the present study, we developed a Massively Parallel Reporter Assay (MPRA) to investigate whether specific 3′ UTR sequences or features influence RNA localization and local translation in astrocytes. MPRAs have emerged as powerful genomics tools for screening effects of non-coding regulatory elements on gene expression^33^ and RNA stability^34,35^, widely in culture and more recently in living brain^33,36^, and mRNA localization in cultured neurons^37,38^. However, *in vitro* approaches have several limitations. First, *in vitro* systems cannot fully replicate dynamic interactions and communication between astrocytes and neurons that occur during development or in response to stimuli, posing a challenge to studying the regulation of genes, such as *Sparc* and *Glt1*, mediating these interactions. Second, because the maturation and complex morphology of astrocytes depends on interactions with neurons and neurotransmitters^39,40^, *in vitro* models fail to fully capture mRNA localization dynamics and translational regulation in PAPs. Indeed, morphological complexity of astrocytes is highly compromised in most culture systems. Therefore, to perform a high-throughput characterization of 3’ UTR elements sufficient to drive *Sparc* and *Glt1a* mRNA localization and local translation in *in vivo* astrocytes, we developed SN-MPRA, which combines the high-throughput capabilities of MPRAs with *in vivo* synaptoneurosome (SN) fractionation and astrocyte-specific translating ribosome affinity purification (TRAP), enabling studies of RNA localization (SN), translation (TRAP), and translation in the SN fraction (PAP-TRAP). We complemented that with a comprehensive mutagenesis of active sequences to define the activity of each element to the nucleotide resolution. Overall, we found evidence for multiple complementary mechanisms governing mRNA localization in astrocytes, including increased abundance and the presence of specific sequence motifs.

## Results

### Development of SN-MPRA

Because *in vivo* MPRAs introduce more challenges to delivery and recovery of sequence elements than cell culture^41^, we designed a focused yet comprehensive ∼6500 element library that covered two locally translated mRNAs in astrocytes: *Sparc* (ENSMUST00000213866.2; Supplemental Figure 1A) and *Glt1a* (*Slc1a2-202;* ENSMUST00000080210.10; Supplemental Figure 1B), a second splice isoform of the same gene with differential localization^42^, *Glt1b* (*Slc1a2-201;* ENSMUST00000005220.11), and one soma-retained gene, *Hsbp1* (ENSMUST00000034300.8)^15^. To determine which *cis-*regulatory sequences within these transcripts drive their localization, we designed a pool of oligos 130 nucleotides (nts) in length, tiled across the 3’ UTRs in 20 nt steps (Figure 1A). Also included were shuffled controls to assess the impact of random sequences and the β-actin zipcode element as a potential positive control (Supplemental Table 1)^27^. Each element was barcoded 10 times. The pool was cloned into the 3’ UTR of a tdTomato reporter downstream of a GFAP promoter and packaged into AAV2/9 (Supplemental Figure 1C).

**Figure 1:**
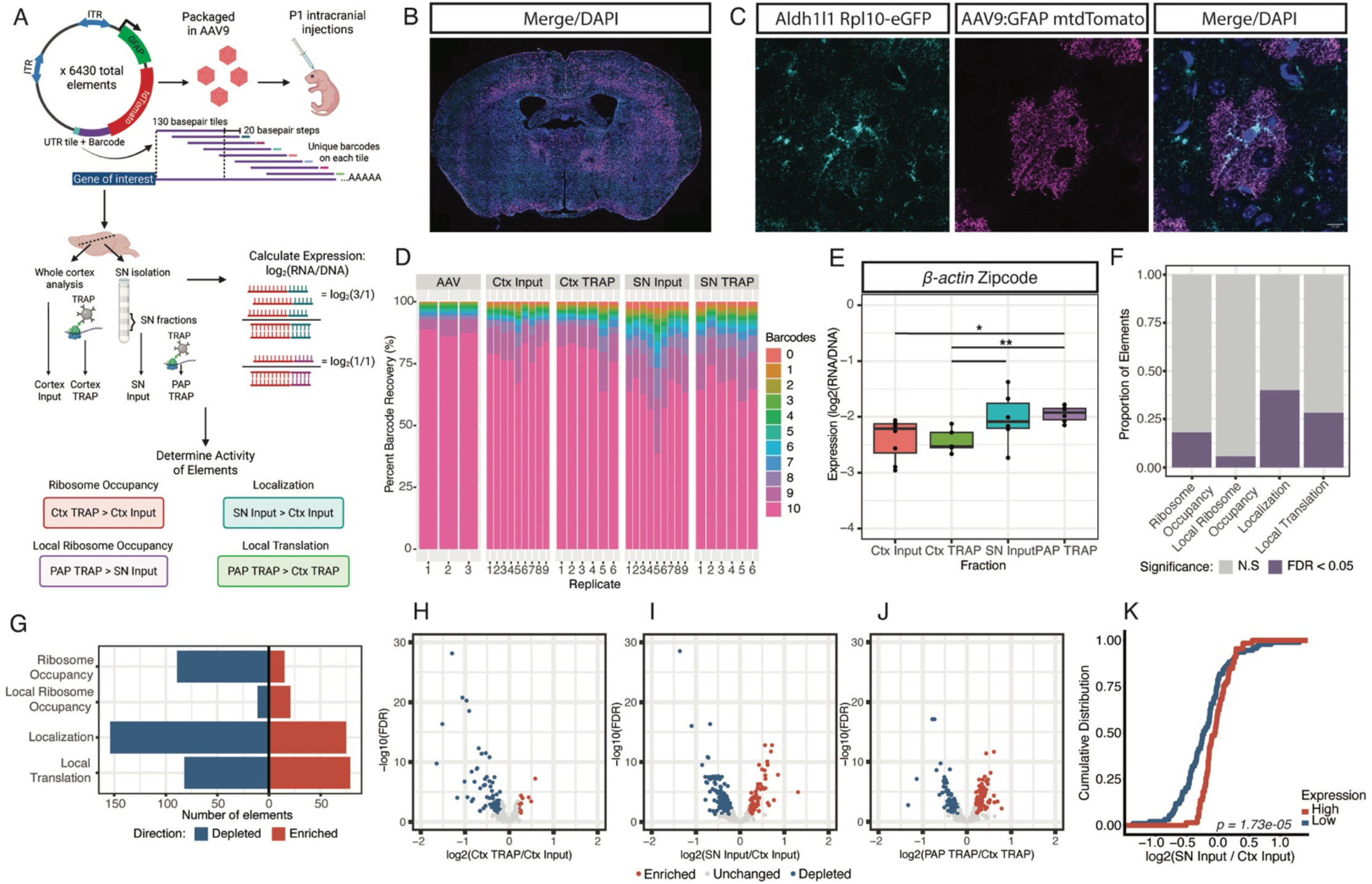
Development of an MPRA Approach for Studying Local Translation *in vivo*. **A**, Schematic of SN-MPRA. (1) Approximately 6430 elements consisting of 130 basepair fragments of 3’ UTRs were tiled in 20 basepair steps. Elements were clone *en masse* in 3’ UTR of tdTomato under a GFAP promoter. The resulting library was packaged into AAV9 and injected into P1 mice. (2) Brains are then harvested for PAP-TRAP. (3) Library specific RNA is isolated and sequenced to enable (3) determination of ribosome occupancy (cortex TRAP vs cortex input), local ribosome occupancy (PAP TRAP vs SN input), localization (SN input vs cortex input), and local translation (PAP TRAP vs cortex TRAP). **B**, Sample coronal slice showing expression of DAPI (blue), tdTomato (magenta) and Aldh1l1:Rpl10a-eGFP (cyan). **C**, Representative astrocyte; scale bar = 10 um. **D**, Distribution of barcode recovery across biological replicates and sample fractions. Stacked bar plots show the percentage of elements with a given number of recovered barcodes (out of 10 total) for each replicate and sample fraction. **E**, Box plot shows the average Expression (log2(RNA/DNA)) of b-actin zipcode control. Significance determined after linear mixed model to account for random effect of barcodes (Expression ∼Fraction + (1|BC)) using pairwise t-test with Tukey’s p-value adjustment. *p<0.05; **p<0.005. **F**. Stacked bar chart showing marginal proportions of Significant (purple) and Not Significant (grey) elements in measures of ribosome occupancy, local ribosome occupancy, localization, and local translation. Significant elements are defined as having an absolute log2FC > 0.2 and an FDR < 0.05. Differences in the proportion of significant elements across fractions were assessed using Cochran’s Q test (Q = 218.11, p-value < 2.2e-16), and paired comparisons were made using McNemar tests with Bonferroni adjustments (RNA localization:ribosome occupancy adjusted p = 8.3e-17; local translation:ribosome occupancy adjusted p = 5.7e-4). **G**. Diverging bar chart showing direction of significance of elements in A (blue = depleted; red = enriched).**H-J**. Volcano plots showing significance (y-axis) vs **H**. ribosome occupancy (log2(cortex TRAP/cortex input)), **I**. localization (log2(SN input/cortex input)), or **J**. local translation (log2(PAP TRAP/cortex TRAP)). Significance determined after linear mixed model to account for random effect of Barcodes (Expression ∼ Fraction + (1|BC)) with FDR correction. Significantly enriched (red) or depleted (blue) elements defined as FDR < 0.05, log2(FC) > 0.25 (enriched) or < -0.25 (depleted). **K**. Cumulative Distribution Function (CDF) shows increased localization (log2(SN Input/Ctx Input)) for elements considered high expression (red) or low expression (blue) based on elements one standard deviation above or below the mean expression of all elements (p = 3.46e-05). Statistical comparisons between distributions were performed using Kolmogorov–Smirnov tests, and corrected for multiple testing with Benjamini Hochberg (BH) FDR adjustment.

To characterize how these sequence elements influence RNA localization *in vivo*, the AAV containing the SN-MPRA library was injected into the cortex of P1 Aldh1l1-eGFP/Rpl10a mice or WT littermates. After validating the AAV transduced cortical astrocytes (Figure 1B,C), we performed PAP-TRAP–an established method that captures astrocyte ribosomes from PAP-enriched synaptoneurosomal fractions^15^ on TRAP+ mice (PAP-TRAP; n = 6). We compared this to astrocyte ribosome-bound reporter mRNAs in both whole cell (via parallel TRAP of cortex; n = 6), RNA from bulk cortex (cortex input; n = 9) and bulk synaptoneurosome (SN input; n = 9) collected from both TRAP+ mice and WT littermates (Figure 1A). From each sample, we quantified the relative abundance of all 6430 candidate regulatory elements using a targeted RNA-sequencing approach. Overall, we achieved robust recovery of barcodes: with typically >70% of barcodes detected (Figure 1D; Supplemental Table 2), with even better recovery from Input and TRAP samples. Likewise, both RNA CPM (counts per million, Supplemental Figure 1D) and RNA/DNA (expression; Supplemental Figure 1E) correlated well across most samples. For downstream analyses, only elements with sufficient coverage were retained, yielding 5,818 total barcodes (∼90% of original library; Figure 1D). Finally, we calculated measures of astrocyte ribosome occupancy (log2(cortex TRAP/cortex input); sometimes called ‘Translational Efficiency’), RNA localization (log2(SN input/cortex input)) and local translation (log2(PAP-TRAP/cortex TRAP)) for each sample.

To validate that our SN-MPRA successfully measured localization, we included the β-actin zipcode^27^ element in our library. Using a linear mixed model to account for the random effects of individual barcodes, we found that the expression of the β-actin zipcode, defined as log2(RNA/DNA), was significantly higher in the PAP TRAP fraction than the cortex input and cortex TRAP (Figure 1E; p = 0.0252 and 0.0012, respectively), indicating that the SN-MPRA design accurately detected differential local translation.

### Tiling SN-MPRA provides insight into regulatory features of 3’ UTRs in astrocytes

We first sought to determine the primary regulatory role of the 3’ UTR generally by comparing the number of elements impacting either ribosome occupancy, localization, or local translation. Historically, while the 5’ UTR is believed to primarily affect translational efficiency^43^, the 3’ UTR is largely thought to influence factors such as RNA stability and localization, with some influence on translational efficiency^44,45^. In line with this, our 3’ UTR elements were more frequently classified as significantly affecting RNA localization and local translation than ribosome occupancy (McNemar’s test, Bonferroni-adjusted p = 8.3e-17, p = 5.7e-4, respectively; Figure 1F). Further, while 15 elements significantly increased ribosomal occupancy, 90 elements significantly decreased it (Figure 1G,H). This directional bias could reflect 3′ UTR–mediated effects on RNA stability that promote degradation and thus secondarily limit translation. Overall, the 3’ UTR tiles had a much more frequent influence on localization and local translation: 84 elements were enriched in measures of localization (Figure 1G,I) and 79 tiles were enriched for local translation (Figure 1G,J). On the other hand, 144 elements were depleted in measures of localization (Figure 1G,I) and 82 were depleted in measures of local translation (Figure 1G,J). It is worth mentioning that very few elements had a significant effect on local ribosome occupancy (Figure 1F,G), which may support a model in which mRNAs are trafficked on stalled ribosomes, as is seen in neuronal RNA granules ^46^. Overall, our results indicate that 3’ UTRs have a higher propensity to affect localization than translational efficiency.

Given that astrocytes do not exhibit the unipolar morphology of neurons, one plausible mechanism of localization is increased transcript abundance to increase the probability of mRNAs reaching the PAPs. Therefore, to determine if localization was largely driven by higher expression, we plotted the Cumulative Distribution of localization values for elements with high or low expression (log2(cortex input RNA/AAV DNA); defined as one standard deviation above or below the mean expression). We found that, while there was a significantly higher localization for elements with high expression (p = 1.73e-05; Figure 1I), there was not a significant change in local translation (Supplemental Figure 2A). Further, although there was a slightly significant correlation between expression and localization for both *Sparc* and *Glt1a* elements (FDR < 0.05; Supplemental Figure 2B,C), this relationship was weak, with several individual elements showing a discordant relationship (i.e. high expression but low localization or vice versa). This suggests that while enhanced expression can increase RNA localization generally, this phenomenon alone does not account for all differences in mRNA localization, motivating us to investigate features of the 3’ UTR elements.

While previous work has identified hundreds of locally translated mRNAs^9,14–17^, relatively few ‘zipcode’ motifs have been characterized^11,27^. However, 3’ UTR features, such as increased GC content and more stable RNA secondary structures, have been found to correlate with RNA localization in various cell types^9,15,17^. We therefore first tested whether ‘sequence content’, such as GC content and predicted structure, or specific sequence motifs were the main drivers of mRNA localization. To do so, we correlated both the GC content and a measure of secondary structures (predicted ΔG) of each element with its activity. We found that while there was a minor but significant correlation with ribosome occupancy (Supplemental Figure 2D,H) and local ribosome occupancy (Supplemental Figure 2E,I), GC content and ΔG did not correlate with mRNA localization (Supplemental Figure 2F,J) or local translation (Supplemental Figure 2G,K). We next looked at the cumulative distributions of our data and similarly found the GC content and ΔG distribution of elements that were enriched in measures of ribosome occupancy were significantly shifted relative to depleted elements, indicating a global difference in GC content and ΔG. Further, when comparing the localization and local translation of the *Sparc* UTR elements and their shuffled controls, we found no correlation (Supplemental Figure 2T,U). Together, these results indicate that sequence content alone is insufficient to explain mRNA localization, motivating us to examine specific sequence motifs.

We therefore examined the activity of each element as it relates to its position in the 3’ UTR. We first determined whether localization and local translation are primarily restricted to biologically derived sequences by comparing native UTR elements to shuffled controls, finding no significant difference between directional localization and local translation (Supplemental Figure 2V-W, respectively) or the magnitude of localization and local translation effect sizes (absolute value of log2FC; Supplemental Figure 2X-Y). This evidence that random sequences can drive a range of localization responses suggests that many possible motifs may promote such activity. Therefore, to find robust *cis*-regulatory candidates sufficient to drive RNA localization, we focused on regions of the 3’ UTRs where 4 or more sequential tiles showed either enriched or depleted localization or local translation, suggesting a shared, singular overlapping motif sufficient to regulate their sub-cellular localization. Consistent with our prior observations, measures of ribosome occupancy showed more frequent depletion across all genes examined, with 3 *Glt1a* element groups displaying consistent ribosome depletion and no element groups displaying consistent ribosome enrichment (Supplemental Figure 3A-E). However, focusing on localization, this analysis identified 8 *Glt1a* element groups (Figure 2A-B) and 2 *Sparc* element groups (Figure 2C) with shared localization. Interestingly *Hsbp1*, a soma*-*retained gene^15^, had no qualifying element groups (Figure 2D). Extending this assessment to local translation, we found 5 *Glt1a* element groups (Figure 2E-F) and 2 *Sparc* element groups (Figure 2G), often overlapping those mediating localization, met this criterion. *Hsbp1* again had no element groups meeting criterion (Figure 2H). These results suggest that, while few motifs can alter ribosome occupancy, shared motifs within our SN-MPRA significantly mediate differential localization and local translation.

**Figure 2:**
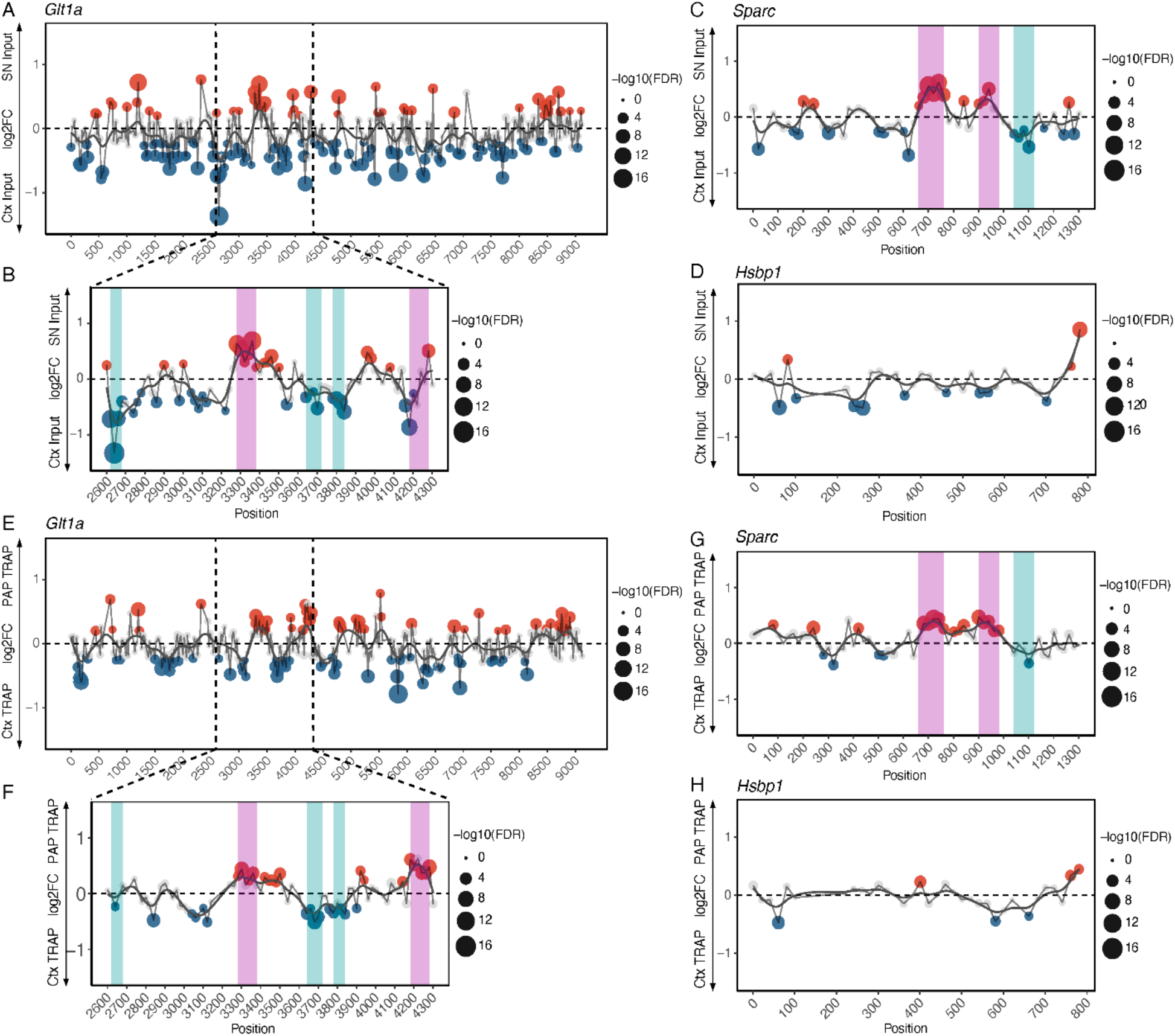
SN-MPRA identifies regions in the 3’ UTRs sufficient to drive RNA localization and local translation. **A-H**, Localization (log2(SN input/cortex input)) or local translation (log2(PAP TRAP/cortex TRAP)) (*y*) of elements plotted against position of tile in the 3’ UTR (*x*). Significance determined after linear mixed model to account for random effect of Barcodes (Expression ∼ Fraction + (1|BC)) with FDR correction. Significantly enriched (red) or depleted (blue) tiles defined as FDR < 0.05, log2(FC) > 0.25 (enriched) or < -0.25 (depleted). Highlighted regions show 4+ consecutive tiles with significant enrichment (magenta) or depletion (cyan) in either localization or local translation measures that were selected for downstream experiments. **A-D**, Localization of *Glt1a* full length 3’ UTR (**A**), including a region with several consecutive effects on localization (**B**), *Sparc* 3’ UTR (**C**), and *Hsbp1* 3’ UTR (**D**) tiles. **E-H**, Local translation of *Glt1a* full length 3’ UTR (**E**), including a region with several consecutive effects on local translation (**F**), *Sparc* 3’ UTR (**G**), and *Hsbp1* 3’ UTR (**H**) tiles.

Our assay also allowed us to investigate differences in activity of two *Glt1* isoforms. Differential localization of these isoforms has been previously characterized, with *Glt1a* mRNA found to be enriched in astrocyte processes and *Glt1b* found to be restricted to the astrocyte soma^42^; however, the factors mediating this are unclear. We therefore compared the activity of both isoforms in our SN-MPRA. Interestingly, while *Glt1a* had tiles that were both significantly soma-retained and significantly localized (Figure 2A-B), the *GLT1b* 3’ UTR tiles had no significantly localized or locally translated tiles (Supplemental Figure 3F-G).

To cost-effectively validate a robust element group, we used postnatal astrocyte labeling by electroporation (PALE)^47^ to probe whether our SN-MPRA was identifying regions of the 3’ UTR necessary and/or sufficient for local translation, focusing on the *Sparc* 3’ UTR which was shorter and thus more experimentally tractable than *Glt1a*. Specifically, the sequence spanning 661-890 showed robust enrichment in measures of both localization and local translation. We designed dual fluorescent constructs containing eGFP with a constant 3’ UTR and mCherry followed by either the *Sparc* 3’ UTR, the *Sparc* 3’ UTR with these nucleotides deleted (*Sparc*Δ661-890) to test necessity, that region of the 3’ UTR alone (*Sparc* 661-890) to test sufficiency, and the *Hsbp1* 3’ UTR (Supplemental Table 4,5). The reporters were flanked by piggybac ITRs, enabling clonal integration into sparse astrocytes when electroporated with a hyperactive piggybac transposase into P0-P1 mice. To determine if these UTRs influenced local translation, we quantified the RFP:GFP ratio as a function of distance from the nucleus using a Sholl-inspired concentric ring analysis (Figure 3A) at P21. Linear mixed models revealed a significant UTR-by-distance interaction, indicating that different UTRs influenced the spatial distribution of RFP. We therefore performed pairwise comparisons at each distance and found that while the *Sparc* 3’ UTR (n = 6 mice; 26 cells) and *Sparc* 661-890 (n = 4 mice; 39 cells) exhibited similar spatial profiles, *Sparc*Δ661-890 (n = 6 mice; 39 cells) had a significantly decreased RFP:GFP ratio in both peri-nuclear and distal regions of the astrocytes (Figure 3B-D,F). Interestingly, the *Hsbp1* 3’ UTR (n = 3 mice; 40 cells) showed minimal RFP signal in all cells (Figure 3D,F). Together, these results indicate that the 661–890 region of the *Sparc* 3′ UTR contributes to robust translation across the cell and is sufficient to alter protein levels at the distal parts of astrocytes. Further, deletion of this region reduces expression at all distances, suggesting this region may play a role in both enhancing overall expression and promoting local enrichment, similar to the CamKIIa 3’UTR element^48^.

**Figure 3:**
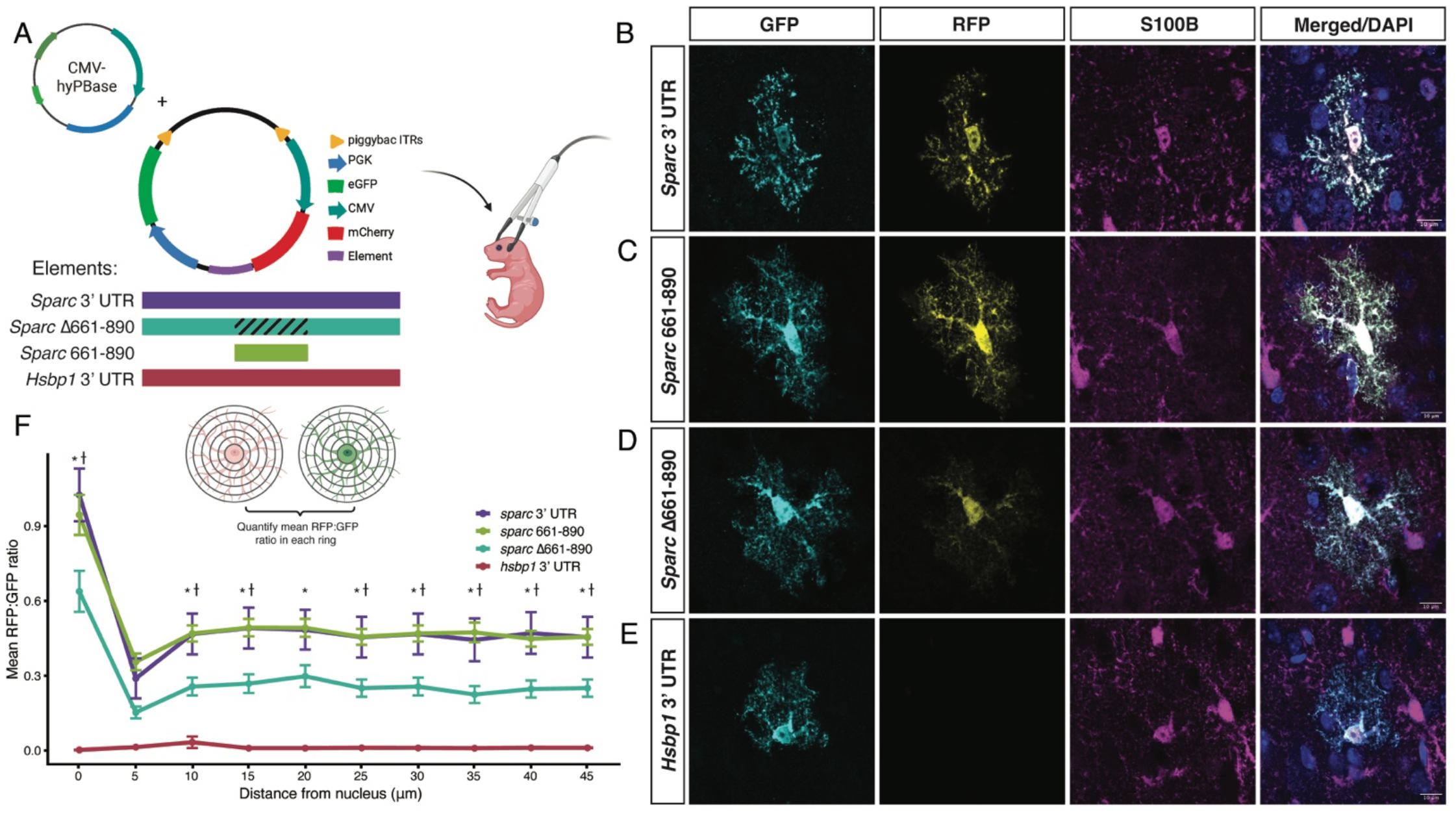
Testing the Sufficiency and Necessity of a Region of the *Sparc* 3’ UTR Identified by the SN-MPRA for Driving Local Translation. **A**. Schematic of constructs and analysis for post-natal electroporations. **B-E**. Representative astrocyte showing GFP (cyan), RFP (yellow), S100B (magenta) and DAPI (blue) for **B**. *Sparc* 3’ UTR, **C**. *Sparc 661-890*, **D**. *Sparc*Δ661-890, and **E**. *Hsbp1* 3’ UTR. (Scale bar = 10 um) **F**. Quantification of RFP:GFP ratio at increasing concentric rings from the nucleus for *Sparc* 3’ UTR (purple), *Sparc* 661-890 (green), *Sparc*Δ661-890 (cyan), and *Hsbp1* 3’ UTR (red). Error bars represent standard error of the mean. Statistical significance was assessed using a linear mixed-effects model (ratio ∼ UTR × distance), with random intercepts for cells nested within mice to account for repeated measurements within cells and animals. This showed a significant effect of UTR (p < 8e-4), distance from nucleus (p < 2.2e-16) and an interaction between UTR and distance from the nucleus (p < 2.2e-16). Pairwise comparisons with BH adjustments were performed for all plasmids at every distance. All comparisons to *Hsbp1* 3’ UTR were significant, but not denoted on the graph. *adjusted p-value < 0.05 (*Sparc* 3’ UTR:*Sparc*Δ661-890); ⍭adjusted p-value < 0.05 (*Sparc 661-890*:*Sparc*Δ661-890).

### Single Nucleotide Mutagenesis SN-MPRA Highlights Sequences That Influence Localization and Local Translation

As the tiling SN-MPRA library enabled us to identify which regions of the 3’ UTR promoted mRNA localization, we next tested which specific nucleotides (and thus motifs) within those regions are necessary for their localization. We generated a second MPRA library containing the center 190-bp of each element group highlighted in Figure 2, including every possible single point mutation, and 20 shuffled controls of each element group. We then repeated PAP-TRAP to determine which nucleotide mutations altered localization and local translation (Figure 4A; Supplemental Table 6). We found robust correlations between CPMs and expression (log2(RNA/DNA)) for most replicates (Supplemental Figure 4A,B), and an overall good distribution of elements packaged in the AAV, with the exception of one element group (*Glt1a* 2621-2681) that did not clone well and was therefore not further analyzed (Supplemental Figure 4C; Supplemental Table 7). A few replicates were found to be significant outliers (Supplemental Figure 4D) and were therefore omitted from downstream analysis.

**Figure 4:**
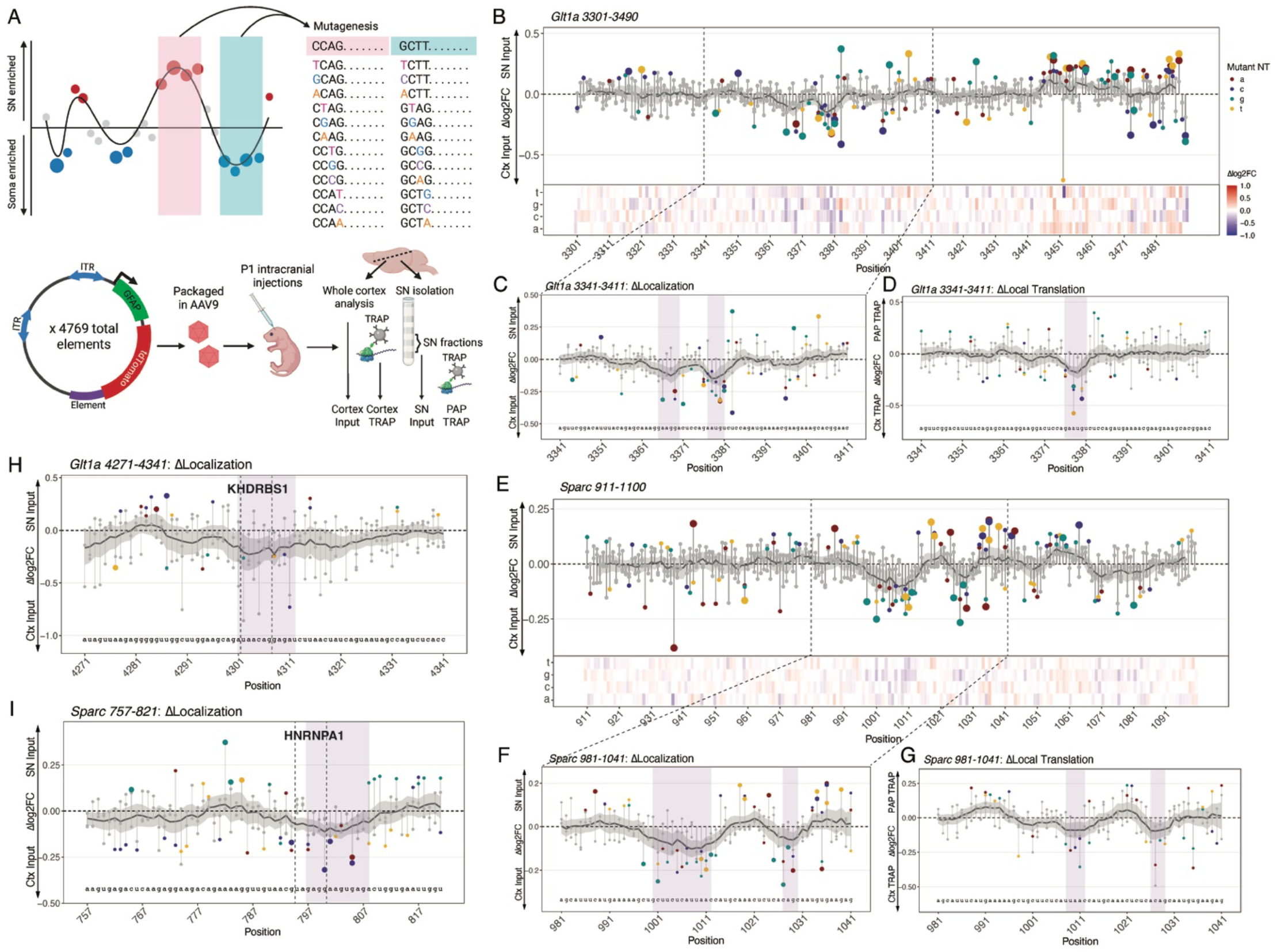
Comprehensive *in vivo* single nucleotide mutagenesis SN-MPRA provides insight into sequence motifs driving localization and local translation. **A**. Schematic of single nucleotide mutagenesis library design. 8 element groups were selected from SN-MPRA results. Consensus sequence for each group was taken as the center 190bps of the overlapping sequences, and the library was designed by mutating every base position to the other three nucleotides. This library was then packaged into AAV9 and delivered to P1 mice via intracranial injections. Brains were harvested for PAP-TRAP at P21. **B**,**E**. ΔLocalization (log2FC(normSN Input/normCtx Input)(*y)* of mutagenesis elements (mutant nucleotide specified by red = A, purple = C, cyan = G, yellow = T/U) plotted against position of individual mutation in the 3’ UTR fragment *(x)* with heat map representation of the Δlog2FC below for **B**. *Glt1a*, positions 3341-3411 and **E**. *Sparc*, positions 911-1100. **C, D**. As in B-E, ΔLocalization **(C)** and ΔLocal Translation **(D)** for a zoomed in portion of *Glt1a*, positions 3341-3411. **F, G**. As in B-E, ΔLocalization **(F)** and ΔLocal Translation **(G)** for a zoomed in portion of *Sparc*, positions 981-1041. **H**,**I**. As in B-G, ΔLocalization for a zoomed in portion of **H**. *Glt1a*, positions 4271-4341, and **I**. *Sparc*, positions 757-821. Dotted lines highlight significant RBPs that overlap regions that disrupt localization. Significant RBP motifs (z ≥ 3) were defined based on enrichment over motif-specific null distributions from randomized sequences. For all measures, significance determined after linear mixed model to account for random effect of replicate (Expression ∼ Fraction + (1|Replicate)) with FDR correction. Grey dots: ns; small colored dots: p < 0.05; larger colored dots: FDR < 0.05. Purple box highlights significant mutation-sensitive regions, defined as contiguous positions with Gaussian-smoothed effects below an adaptive threshold. Significance determined using a one-sided Wilcoxon test comparing mutations within versus outside each region with FDR correction.

To quantify the effects of individual mutagenized elements, we calculated Δribosome occupancy (log2(normCtx-TRAP/normCtx Input); Supplemental Figure 5A-G), Δlocalization (log2(normSN Input/normCtx Input); Figure 4B,C,E,F,H,I; Supplemental Figure 6A-E), and Δlocal translation (log2(normPAP-TRAP/normCtx-TRAP); Figure 4D,G; Supplemental Figure 7A-G; full results Table 8). While 3’ UTR elements originally had minimal effects on ribosome occupancy, mutations to motifs such as CUGUCUG (Supplemental Figure 5C), polypyrimidine tracts (Supplemental Figure 5D), or an upstream poly-A signal (Supplemental Figure 5F) led to increased ribosome occupancy. Further, these analyses revealed regions within previously localized elements where nucleotide substitutions consistently reduced localization (Figure 4B,C,E,F,H,I; full-length elements Supplemental Figure 6), with a few corresponding regions disrupting local translation (Figure 4D,G; full-length elements Supplemental Figure 7). Interestingly, while some regions overlapped, the nucleotides disrupting localization did not uniformly affect local translation (Figure 4B-G), potentially indicating mechanistic divergence between these processes. Notably, element groups that were significantly depleted from the synaptoneurosome fractions in the first library, such as *Sparc* 1051-1240, appeared to have the strongest disruption from the mutagenesis. These findings highlight specific sequence regions that are important for localization and, in some cases, local translation.

As our tiling library showed that increased expression could partially explain differential localization of some elements, we next asked whether mutagenesis elements changing expression were similarly associated with changes in localization by correlating Δexpression with Δlocalization for each element group (Supplemental Figure 8A-G). We applied stringent significance and effect-size criteria (Spearman rho > 0.25, FDR < 0.01) to identify robust associations. Using this approach, we identified three element groups in which Δexpression and Δlocalization were significantly correlated, suggesting that processes influencing expression, such as transcription or RNA stability, may also enable differential localization.

It is well established that 3’ UTRs of locally translated mRNAs in the brain are longer, more conserved, and form more stable secondary structures^9,15,44,49^. Among secondary structures, G-quadruplexes have been shown to bind FMRP and promote mRNA localization in neurons ^30^. We therefore asked whether changes in ribosome occupancy, localization, or local translation could be predicted by sequence conservation, and similarly whether these measures significantly differ from mutations that result in changes to secondary structures (henceforth referred to as ΔΔG) or propensity to form G-quadruplexes (referred to as ΔrG4; results summary Supplemental Table 9)^50^. Overall, ΔΔG showed no discernible relationship with Δribosome occupancy (Supplemental Figure 9A) or Δlocal translation (Supplemental Table 9), and weak correlation with Δlocalization in one element group (Supplemental Figure 9C). Notably, this association indicated that increases in ΔΔG, and therefore decreased secondary structure stability, modestly correlated with increases in Δlocalization. Similarly, conservation scores were largely not-predictive, correlating with Δribosome occupancy in only a single element group, but showing no relationship with Δlocalization and Δlocal translation (Supplemental Table 9). Finally, ΔrG4 did not correlate with Δlocal translation, but exhibited a weak inverse correlation with Δribosome occupancy in one element group (Supplemental Figure 9B) and with Δlocalization in two element groups (Supplemental Figure 9D). Although our library was not designed to intentionally include G-quadruplexes, and ΔrG4 reflects only propensity rather than confirmed structure, these results were nonetheless unexpected.

We next investigated whether the sequences disrupting localization also disrupted RBP motifs by aligning our results to the ATtRACT (A daTabase of RNA binding proteins and AssoCiated moTif) motif database^51^. Focusing on element groups that were previously localized, we found overlap between RBP motifs such as KHDRBS1 (Figure 4H) and HNRNPA1 (Figure 4I) with regions of our mutagenesis library that disrupted localization. Interestingly, KHDRBS1, or *Sam68*, has previously been found to regulate post-synaptic β*-*actin metabolism ^52^, suggesting this RBP may be a mediator of *Glt1a* localization. Element groups that were significantly depleted from the SN fractions also had RBP motifs, such as HNRNPF, HNRNPH1, and RBM38, corresponding to sequences further disrupting localization (Supplemental Table 10). However, not all regions that disrupt localization correspond to a known RBP motif. Together, our results suggest that RNA localization in astrocytes is likely mediated by multiple mechanisms, rather than by classical “zipcode” motifs alone (Table 1).

**Table 1:** Results Summary.

| Element group | Expression | G-quadruplex | $\Delta G$<br>(kcal/mol) | RBP Motif |
| --- | --- | --- | --- | --- |
| Glt1a<br>3301-3490 |  |  | ✓ |  |
| Glt1a<br>3641-3830 |  |  |  | ADARB1 |
| Glt1a<br>3781-3970 |  |  |  | RAVER1 |
| Glt1a<br>4201-4390 | ✓ | ✓ |  | SRSF1/LIN28A/KHDRBS1 |
| Sparc<br>681-870 |  | ✓ |  | HNRNPA1 |
| Sparc<br>911-1100 | ✓ |  |  |  |
| Sparc<br>1051-1240 | ✓ |  |  | RBM38 |

Table summarizing the different mechanisms (expression, propensity to form G-quadruplexes, measures of secondary structure, and presence of RBP motifs in disrupted regions) underlying changes in localization based on mutagenesis SN-MPRA results. Check marks describe element groups that had significant relationships in each analysis.

## Discussion

An asymmetric distribution of mRNAs in cells can be achieved by several complementary mechanisms: on one hand, there can be very precise mechanisms involving selective targeting via RBP interactions with *cis-* regulatory features such as zipcode motifs ^27–29^ or secondary structures ^30^. Alternative mechanisms may include enhancing the overall expression or stability to increase probability of mRNAs arriving at the periphery (or, alternatively, enhanced degradation of mRNAs that are soma-retained). To understand which of these possibilities apply in astrocytes, we developed an *in vivo* MPRA to identify which sequences or features in the 3’ UTRs of two locally translated genes, *Sparc* and *Glt1a*, are sufficient to drive mRNA localization and local translation. By combining the high-throughput capabilities of MPRAs with sub-cellular fractionation and astrocyte-specific TRAP, we were able to uncover several regions in the 3’ UTR of these genes that significantly influence localization and local translation. Further, our data provide novel insights into the regulatory roles of 3’ UTRs in astrocytes.

The 3’ UTR is largely believed to influence factors such as RNA stability and sub-cellular localization. In line with this, we found more significant effects on localization and local translation than ribosome occupancy (Figure 1F-H). Given that most of the effects on ribosome occupancy caused a depletion from ribosomes, we hypothesize this is secondary to decreasing RNA stability. Locally translated mRNAs have been found to have higher expression and more stable secondary structures in multiple cell types^9,15^, prompting us to test whether the differential localization was driven by increased expression, 3’ UTR features such as stability or GC content, or specific sequence motifs. Surprisingly, more stable secondary structures or higher GC content did not correspond to an increase in localization or local translation but did have a mild correlation with ribosome occupancy. While there was also a significant association between expression and localization, arguing this can be a mechanism to globally drive local translation, several elements showed a discordant relationship between the two measures suggesting expression alone does not explain all differential localization.

We therefore focused on eight regions of the 3’ UTRs that had at least four consecutive elements driving localization. One such element group in the *Sparc* 3’ UTR, which encompassed tiles spanning from nucleotides 661-890, showed significant enrichment in measures of both localization and local translation. Cloning this fragment into a dual fluorescence reporter, as well as the full *Sparc* 3’ UTR and *Sparc*Δ661-890, revealed that this 3’ UTR region enhanced mCherry protein expression across the entire astrocyte, suggesting this region may influence gene expression levels or mRNA stability, thus also increasing relative protein levels in PAPs. Further, single nucleotide mutagenesis of the eight element groups revealed several sequence regions in which mutations disrupted mRNA localization and local translation. While few of these regions overlapped with known RBP motifs, such as KHDRBS1–previously shown to regulate post-synaptic β*-*actin ^52^, several regions did not correspond to a known RBP motif. This could suggest several mechanisms of mRNA localization in astrocytes exist, including but not limited to mRNA:RBP interactions, increased expression, and increased mRNA stability.

There are a few limitations of this study that should be addressed. First, because the tiling library was 130 nt fragments, localization mediated by large secondary structures, long zipcodes–such as the dendritic targeting element in *Arc* mRNA^53^–or long-range interactions may have been missed. Second, while *in vitro* studies of axon localized mRNAs^38^ had large effect sizes, our studies showed mild enrichment of elements in the SN fractions. This is largely due to the challenges of *in vivo* fractionation and the differences in comparisons. For example, in cultured neurons, it is possible to completely isolate the soma from the neurites, so a direct comparison can be made between the two sub-compartments. *In vivo*, however, SN isolation only enables an enrichment of fine processes, rather than a perfect purification. Additionally, instead of a direct comparison between the SN fractions and soma, we have to make a comparison between the SN fractions and whole cell input, which further reduces effect sizes. Despite these limitations, we were able to characterize sequences important for RNA localization in astrocytes. Further, our system had the clear advantage of studying astrocytes in their native biological context, as the complex morphology of astrocytes is not fully recapitulated *in vitro*^39,40,54^. Indeed, even neurons may benefit from adapting an SN-MPRA to understand their *in vivo* RNA localization, as it may differ substantially from *in vitro* models. Of course, MPRA libraries *in vivo* need to be less complex than those *in vitro*^41^, necessitating our focus on libraries of <10,000 elements here.

Finally, future studies with our SN-MPRA methodology can be used to address many on-going questions in the field. For example, designing a library that contains 3’ UTR tiles from genes in a shared pathway may help uncover whether there is a shared regulatory logic mediating their localization, or if regulation is largely gene specific. Further, combining this with methods to elicit neuronal activity, such as a seizure induction^24^, behavioral manipulations, or chemogenetics, could help define which *cis-*regulatory sequences are important for mediating astrocytic local translational responses to neuronal activity. Lastly, as technology for oligo pool synthesis continues to advance, the use of longer 3’ UTR fragments may enable high-throughput studies of motif necessity or testing of combinatorial logic, such as that of the *MBP* mRNA^11^.

## Supporting information

Combined Supplemental Tables

## Acknowledgments

We want to thank Colin Florian for help with computational pipeline, Mike Vasek for technical support, members of the Cohen, Dougherty, Miller, Zaher, and Djuranovic labs for helpful discussions, and Mingjie Li of the Hope Center Viral vectors core for packaging. Funding was provided by R01NS102272, R01MH116999, and T32GM139774.

## Materials and Methods

### Animals

All procedures were approved by the Institutional Animal Care and Use Committee at Washington University in St. Louis. Veterinary care and housing was provided by the veterinarians and veterinary technicians of Washington University School of Medicine under Dougherty lab’s approved IACUC protocol. TRAP protocols were completed with *FVB-Tg(Aldh1l1-EGFP/Rpl10a)JD130Htz/J* (RRID:IMSR_JAX:030247, The Jackson Laboratory).

### MPRA Library Plasmid Preparation

To generate the tiling library sequences, the 3’ UTR of known astrocyte genes from a previously published dataset ^15^ were segmented into 130 nt tiles with a 20 nt shift and barcoded 10 times with a unique 9 nt identifier. A pool of the corresponding oligos (Table 1) flanked by adapter and restriction site sequences was synthesized by Agilent. Elements were PCR-amplified and inserted into an adeno-associated virus (AAV) plasmid containing the GFAP promoter driving tdTomato in the 3’ UTR position and packaged into AAV9 by the Hope Center Viral Vectors Core.

To generate the single nucleotide mutagenesis library, we chose 3’ UTR fragments where at least 4 consecutive tiles were all either enriched or depleted from the localization or local translation analyses, yielding 8 fragments: *sparc_661_761, sparc_901_981, sparc_1041_1121, Glt1a_2621_2681, Glt1a_3281_3381, Glt1a_3641_3721, Glt1a_3781_3841*, and *Glt1a_4181_4281*. The consensus sequence was generated as the center 190 nts of the overlapping tiles. For each fragment, we then generated each potential base substitution as the mutagenesis tiles, as well as 20 scrambled controls per fragment. A pool of the corresponding oligos (Supplementary Table 6) flanked by adapter and restriction site sequences were synthesized as an oligo pool from Twist Biosciences. Elements were PCR-amplified and cloned in the same backbone as the tiling library. Plasmids were packaged into AAV9 by AAVnerGene.

### *In vivo* SN-MPRA

The AAV9 library was intracranially injected into P1 Aldh1l1-eGFP/Rpl10a litters. At P21, brains were harvested and the cortex dissected for PAP TRAP as described ^15^. Briefly, 3 cortices were pooled per replicate and homogenized in 3 mL ice-cold homogenization buffer using a 7 mL glass dounce homogenizer. Homogenate was spun at 1000 x g for 10 min at 4°C to pellet out nuclei and cell debris. The supernatant was split into two samples: for the input and astrocyte TRAP samples, 500μL was incubated with 50 μL of salt lysis buffer and 50 μL DHPC for 15 minutes on ice and spun at 20,000 x g for 15 min. 60 μL was removed for whole cortex input, and TRAP was performed on the remaining supernatant. For the SN input and PAP TRAP samples: the remaining supernatant (∼2 mL) was layered over a sucrose-Percoll gradient and spun at 32,500 x g for 5 min as described ^55^. The SN fractions were collected by puncturing the bottom of the tube. The SN fractions were diluted with homogenization buffer and spun at 7000 x g for 20 min to pellet SNs and remove Percoll. The supernatant was removed and the pellet was resuspended in 1 mL salt lysis buffer and 100 μL DHPC. 200 μL was removed for SN input, and TRAP was performed on the remaining sample. TRAP samples were incubated at 4°C for 3-4 hours with end over end rotation. After incubation, TRAP samples were washed 4 times with a high salt wash buffer and resuspended in 250 μL wash buffer. RNA was extracted using Zymo Quick-RNA microprep kit (Cat #R1050) or Qiagen RNeasy Mini kit (Cat #74104).

### MPRA Sequencing Library Preparation

Libraries were prepared by taking extracted RNA and performing cDNA synthesis using Superscript IV Reverse Transcriptase standard protocol with library specific priming (GGCACTGGAGTGGCAACT). Resulting cDNA or AAV DNA were then PCR amplified using NEBNext Ultra II Q5 2x Master Mix and library specific forward (CTGACAGGCGCGCCCGAGCTGTACAAGTAAGTCGAC) and reverse (CTGCTCGAGGCAAGCTT) primers, with the forward primer adding an AscI restriction site. Reactions were purified using Magbind beads (Omega Bio-Tek; M1378-00) between each step. The purified PCR products were digested with AscI and HindIII restriction enzymes for 1h at 37°C. The purified digested products were ligated to 4 equimolar staggered adapters, purified, and used for PCR with Illumina primers for library indexing. The purified libraries were then subjected to quality control before 2 × 150 next generation sequencing on an Illumina NovaSeq.

### MPRA Libraries: Barcode and Element Counting

Paired end read files were merged using PANDAseq. Merged sequences were then aligned to the barcodes in the oligonucleotide library using a custom counting script designed to identify exact matches along the entire sequence. For the mutagenesis library, the custom counting script was designed to identify exact matches of the entire element + restriction sites instead of barcodes. Count files were then imported into R for downstream analyses.

### MPRA Libraries: Statistical Analyses

For the tiling library, barcoded elements were CPM normalized before filtering. Barcodes with fewer than 20 counts and elements with fewer than 7 sufficient barcodes were excluded from analysis, yielding a final library of 5,818 sequences. All analyses were conducted with putative outlier samples both included and excluded, and results were largely concordant. Thus, we included them here.

For the mutagenesis library, elements were CPM normalized, and the element group with low expression in all fractions was excluded from downstream analyses.

For both libraries, pairwise CPM and Expression Spearman correlations were performed using the cor() function in R. Replicate/Fraction outliers were identified based on PCA distances from group centroid based on variance stabilizing transformation of DESeq2 object.

For the tiling library, significance in measures of ribosome occupancy (log2(Ctx-TRAP/Ctx Input)), localization (log2(SN Input/Ctx Input)), local translation (log2(PAP TRAP/Ctx TRAP), local ribosome occupancy (log2(PAP TRAP/SN Input)) were determined using a linear mixed model, with barcode as a random effect (Expression ∼ Fraction + (1|BC)) and FDR correction (Benjamini Hochberg procedure). Significant elements were defined as having an absolute log2FC > 0.2 and an FDR < 0.05.

For the mutagenesis library, individual element CPMs were normalized to the median CPM of all elements within that element group before calculating Δribosome occupancy (log2(normCtx-TRAP/normCtx Input)), Δlocalization (log2(normSN Input/normCtx Input)), and Δlocal translation (log2(normPAP-TRAP/normCtx-TRAP)). Significance was determined using a linear mixed model with replicate as a random effect (Normalized Expression ∼ Fraction + (1|Replicate)).

### RNA Binding Protein Motif Discovery

Regions with disrupting mutations were identified by comparing the log2FC of all single-nucleotide variants to wildtype sequences for each element. Mutation effects were summarized at each position (median log2FC), smoothed using a Gaussian sliding window, and positions with smoothed effects below a combined threshold were flagged. Contiguous runs of at least three positions below threshold were called candidate regions, expanded to neighboring positions until signal recovery, merged if overlapping, and rescored by comparing mutations within versus outside the region using a one-sided Wilcoxon rank-sum test with BH correction.

To identify RBP motifs corresponding to disrupting regions, the sequences corresponding to the flagged regions plus the flanking 7 nucleotides on either end were scanned for RBP binding motifs using *Mus musculus* position weight matrices from the ATtRACT database^51^, retaining one representative motif per RBP based on maximal per-position information content. Motif matches were scored using log-odds matrices in a sliding-window framework, and motif-specific null score distributions were generated from randomized sequences to account for differences in motif length and composition. Significant motif occurrences were defined as sites with scores ≥3 standard deviations above the empirical null distribution (z ≥ 3) that overlapped the disrupted region.

### Dual Fluorescence Reporters

The dual fluorescence reporter plasmid combined a CMV enhancer/promoter driving a mCherry2 ORF with an hGH transcription termination signal with a PGK promoter driving an eGFP ORF with a SV40 transcription termination signal. The mCherry2 portion contains NheI/KpnI sites in the 3’ UTR to clone in elements. Validation constructs (Table 4) were cloned into these sites. For validation constructs, *Sparc* 661-890 and *Hsbp1* 3’ UTR were ordered as gene blocks from Twist Biosciences. *Sparc* 3’ UTR was amplified from an existing plasmid ^15^. *Sparc*Δ661-890 was cloned using PCR stitching (primers in Table 5) with Phusion High-Fidelity PCR Master Mix (NEB, M0531). To clone constructs into the reporter plasmid, the plasmid and validation constructs were separately digested with KpnI-HF (NEB, R3142L), NheI-HF (NEB, R3131) in rCutsmart buffer (NEB, B6004) and gel purified with NucleoSpin Gel & PCR Clean-up Mini Kit (Macherey-Nagal, 740609.50). Constructs were ligated into plasmid with T4 DNA Ligase (NEB, M0202). All plasmids were verified by sequencing through Plasmidsaurus.

### Postnatal Astrocyte Labeling by Electroporation (PALE)

Dual fluorescence reporters were delivered to astrocytes based on a previously published protocol ^47^. Briefly, a final mass of 2μg in 1μL of an equimolar mix of CMV-hypBACase and PGK-eGFP-CMV-RFP containing was injected into the lateral ventricle of P0 CD1 mice, and then electroporated with 5x 100 V/50 ms pulses at a 950 ms interpulse interval using ECM 830 Electro Square Porator with 5mm Platinum Plated Tweezertrodes (BTX Harvard Apparatus), with the positive electrode placed above the cortex on the side of injection.

### Dual Fluorescence Reporters

Mice were perfused with ice-cold 1x PBS followed by 4% paraformaldehyde (PFA). Brains were harvested and postfixed overnight in 4% PFA, transferred to 15% sucrose solution for 24 h, then to 30% sucrose solution for another 24 h, frozen in OCT compound (Thermo Fisher, 23-730-571). Brains were then cryosectioned to 30 μM (Leica CM1950). Sections were blocked in blocking buffer (5% Normal Donkey Serum (Jackson Immunoresearch) in 1X PBS + 0.1% Triton-X) for 1h at room temperature. Primary antibodies were incubated at 4°C overnight in blocking buffer (Chicken @ GFP, 1:1000 (Aves, GFP-1020); Rabbit @ RFP, 1:1000 (Rockland, 600-401-379); Guinea Pig @ S100B, 1:250 (Synaptic Systems, 287004)). Alexa Fluor secondary antibodies (Donkey @ Chicken 488, 1:1000 (Jackson ImmunoResearch, 703-545-155); Donkey @ Rabbit 568, 1:1000 (Invitrogen, A10042) Donkey @ Guinea Pig 647, 1:1000 (Jackson ImmunoResearch, 706-605-148)) were incubated for 1 h at room temperature in blocking buffer. Nuclei were counterstained with DAPI at a 1:20,000 dilution, washed, then mounted with Prolong Gold anti-fade mounting medium (Thermo Fisher, P36934). Confocal microscopy was performed on a STELLARIS 5 Confocal Microscope (Leica) and slide scanned images were gathered on Leica Thunder DMi8.

### Code and Data Availability Statement

Code are available on Github at: https://github.com/Dougherty-Lab/astrocyte_sn-mpra. Raw sequencing data being deposited at GEO: (accession pending). Processed data are found in supplemental tables 1-10.

## Supplemental Figures

**Supplemental Figure S1:**
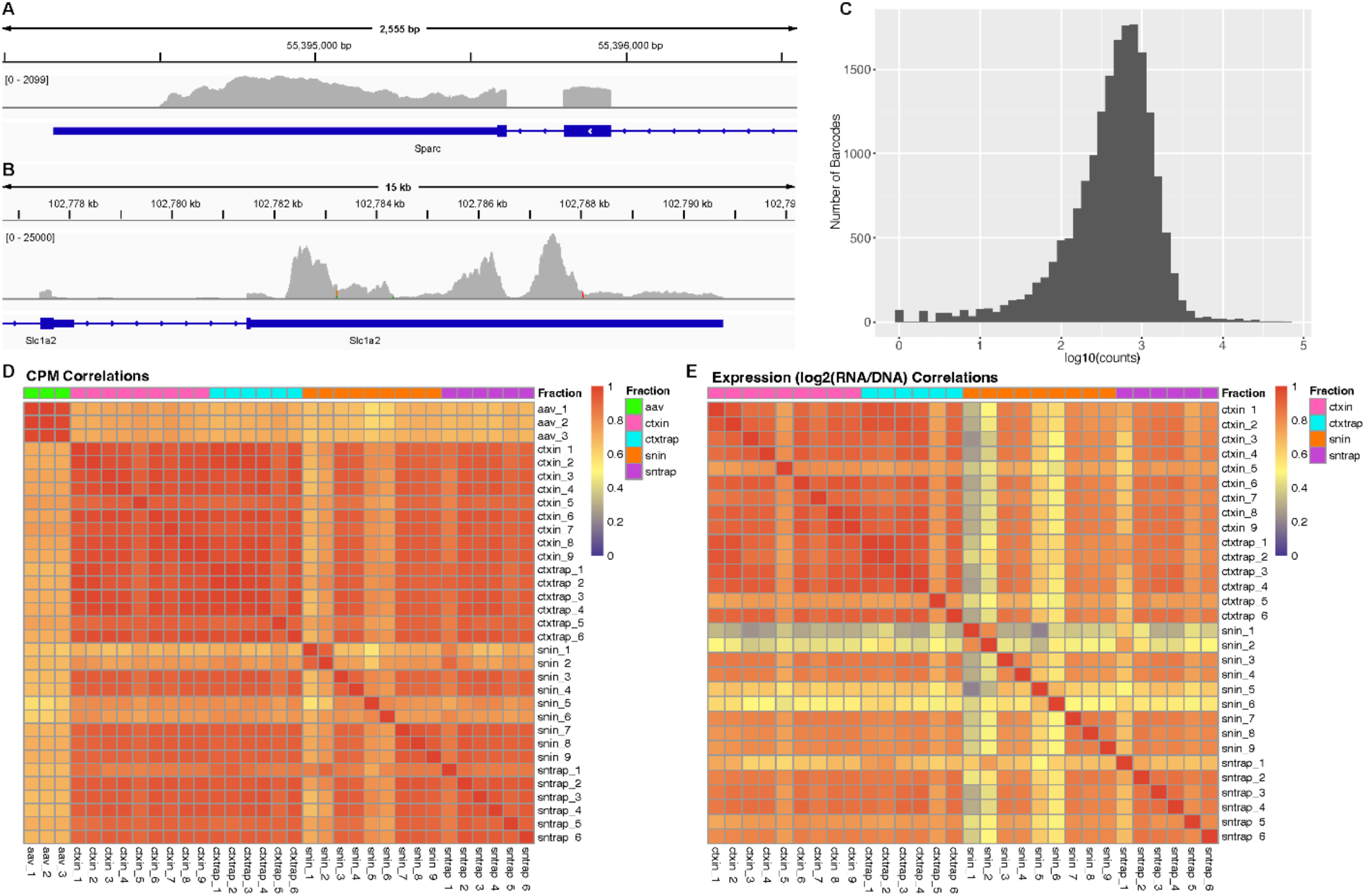
SN-MPRA Library Design and QC Metrics. **A-B**, Integrated Genome Viewer tracks showing PAP TRAP reads aligning to the chosen isoforms of *Sparc* (A; ENSMUST00000213866.2) and *Glt1a* (*Slc1a2-202*) (B; ENSMUST00000080210.10) ^15^ . **C**, Spread of element distribution (log10(counts)) recovered AAV DNA (n=3). Elements that were not cloned were removed for downstream analyses. **D-E**, Heatmaps showing Spearman correlations of (D) CPM and (E) Expression (log2(DNA/RNA)) between replicates and fractions.

**Supplemental Figure S2:**
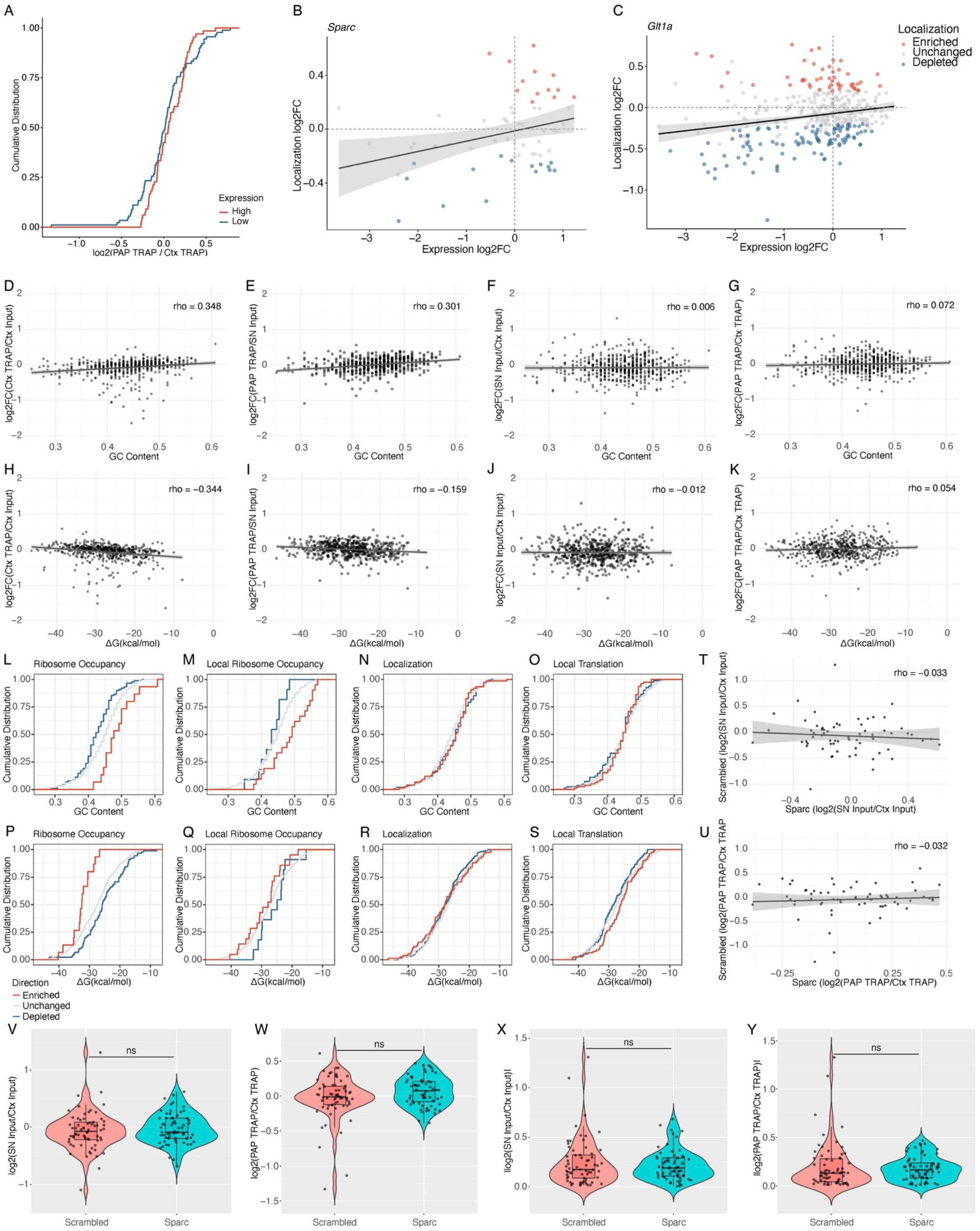
Effects of Expression, GC Content, and RNA Stability on Ribosome Occupancy, Localization, Local Translation, and Local Ribosome Occupancy. **A**. Cumulative Distribution Function (CDF) shows no change in local translation (log2(PAP TRAP/cortex TRAP)) for elements considered high expression (red) or low expression (blue) based on elements one standard deviation above or below the mean expression of all elements (FDR = 0.18). Statistical comparisons between distributions were performed using Kolmogorov–Smirnov tests and corrected for multiple testing with BH FDR correction. **B, C**. Spearman correlations of expression (log2(cortex input/AAV DNA)) and localization (log2(SN input/cortex input)) shows a significant but mild correlation for **B**. *Sparc* 3’ UTR elements (rho = 0.25; p = 0.045) and **C**. *Glt1a* 3’ UTR elements (rho = 0.19; p = 0.0001). Dots colored based on significantly enriched (red), depleted (blue), or unchanged (gray) for measures of localization. **D-G**. Spearman correlations of each element’s GC content with its **D**. ribosome occupancy (log2(cortex TRAP/cortex input); rho = 0.348, p < 2.2e-16), **E**. local ribosome occupancy (log2(PAP TRAP/SN Input); rho = 0.301, p = 2.32e-13), **F**. localization (log2(SN input/cortex input); rho = -0.006, p = 0.894), or **G**. local translation (log2(PAP TRAP/cortex TRAP); rho = 0.072, p = .084)). **H-K**. Spearman correlations of each element’s ΔG with its **H**. ribosome occupancy (log2(cortex TRAP/cortex input); rho = -0.344, p < 2.2e-16 ), **I**. local ribosome occupancy (log2(PAP TRAP/SN input); rho = 0.159, p = 0.0001), **J**. localization (log2(SN Input/cortex input); rho = -0.012, p = 0.776), or **K**. local translation (log2(PAP TRAP/cortex TRAP); rho = 0.054, p = 0.2)). **L-S**. Cumulative distribution functions (CDFs) of **L-O**. GC content and **P-S**. ΔG are shown for elements that significantly increased (up-regulated; [red]), significantly decreased (down-regulated; [blue]), or showed no significant change(unchanged; [grey]) in **L**,**P**. ribosome occupancy (**L**. GC content, D = 0.51, FDR = 0.003; **P**. ΔG, D = 0.60, FDR = 0.0003); **M**,**Q**. local ribosome occupancy (**M**. GC content, D = 0.52, FDR = 0.041; **Q**. ΔG, D = 0.31, FDR = 0.479); **N**,**R**. localization (**N**. GC content, D = 0.09, FDR = 0.82; **R**. ΔG, D= 0.10, FDR = 0.712); or **O**,**S**. local translation (**O**. GC content, D = 0.12, FDR = 0.609; **P**. ΔG, D = 0.19, FDR = 0.15). Statistical comparisons between distributions were performed using Kolmogorov–Smirnov tests and corrected for multiple testing with BH FDR correction. **T**,**U**. Spearman correlations of **T**. localization (rho = -0.033, p = 0.800) and **U**. local translation (rho = -0.032, p = 0.804) for *Sparc* 3’ UTR elements and their shuffled controls. **V-Y**. Violin plot shows the distribution of **V**. directional localization activity (log2FC(SN input/cortex input); p = 0.39), **W**. directional local translation activity (log2FC(PAP TRAP/cortex TRAP); p = 0.14). **X**. magnitude of localization effects (|log2FC(SN input/cortex input)|; p = 0.92), and **Y**. magnitude of local translation effects (|log2FC(PAP TRAP/cortex TRAP)|; p = 0.43). Box plots show median values with interquartile range. Significance determined using Wilcoxon rank sum test.

**Supplemental Figure S3:**
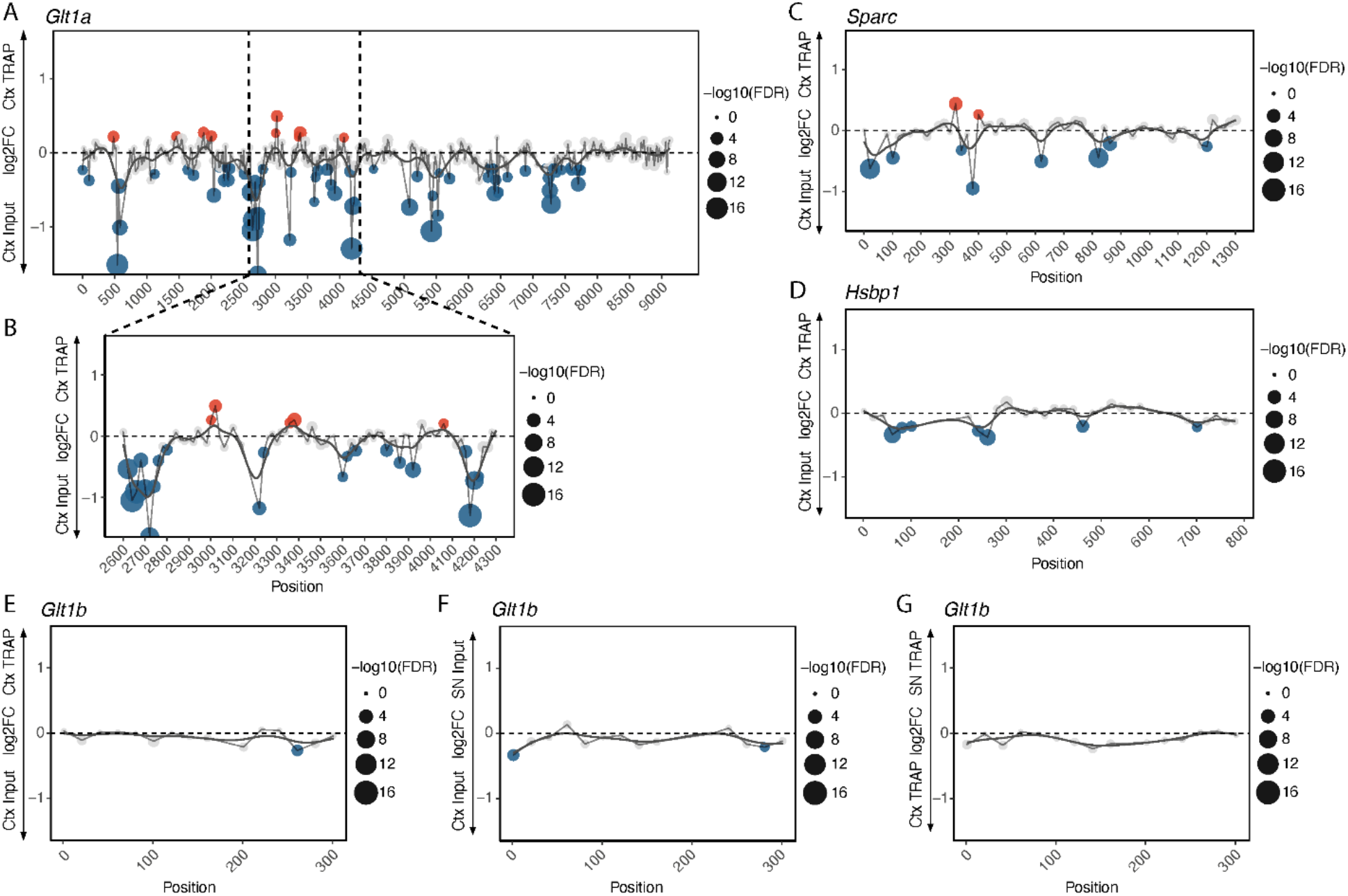
SN-MPRA Extended Results. **A-D**, Ribosome occupancy (log2(cortex TRAP/cortex input)) (*y*) of *Glt1a* full length 3’ UTR (**A**), including the region highlighted in the localization analysis (**B**), *Sparc* 3’ UTR (**C**), and *Hsbp1* 3’ UTR (**D**) tiles plotted against position of tile in the 3’ UTR (*x*). **E-G**, Ribosome occupancy (**E**), localization (**F**), or local translation (*y*) of *Glt1b* tiles (*x*). Significance for **A-G** determined after linear mixed model to account for random effect of Barcodes (Expression ∼ Fraction + (1|BC)) with FDR correction. Significantly enriched (red) or depleted (blue) tiles defined as FDR < 0.05, log2(FC) > 0.25 (enriched) or < -0.25 (depleted).

**Supplemental Figure S4:**
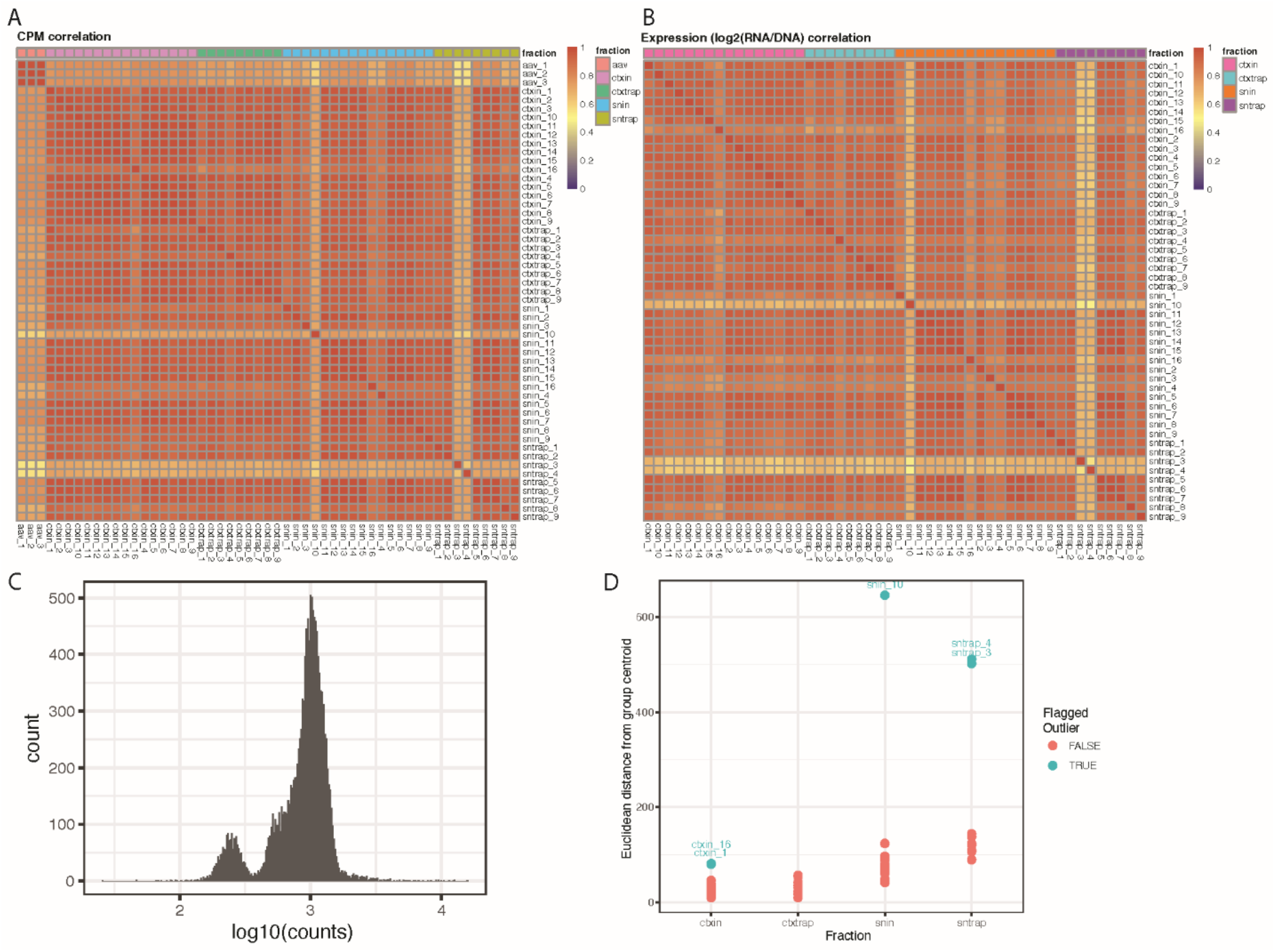
Single Nucleotide Mutagenesis SN-MPRA QC Metrics. **A**,**B**. Heatmaps showing Spearman correlations of (D) CPM and (E) Expression (log2(DNA/RNA)) between replicates and fractions. **C**. Spread of element distribution (log10(counts)) recovered AAV DNA (n=3). One element group did not clone efficiently (left peak) and was removed for downstream analyses. **D**. Technical outliers were flagged and removed based on Euclidean distances of each sample from its fraction-specific centroid in PCA space: samples exceeding median + 3×MAD were flagged and removed.

**Supplemental Figure S5:**
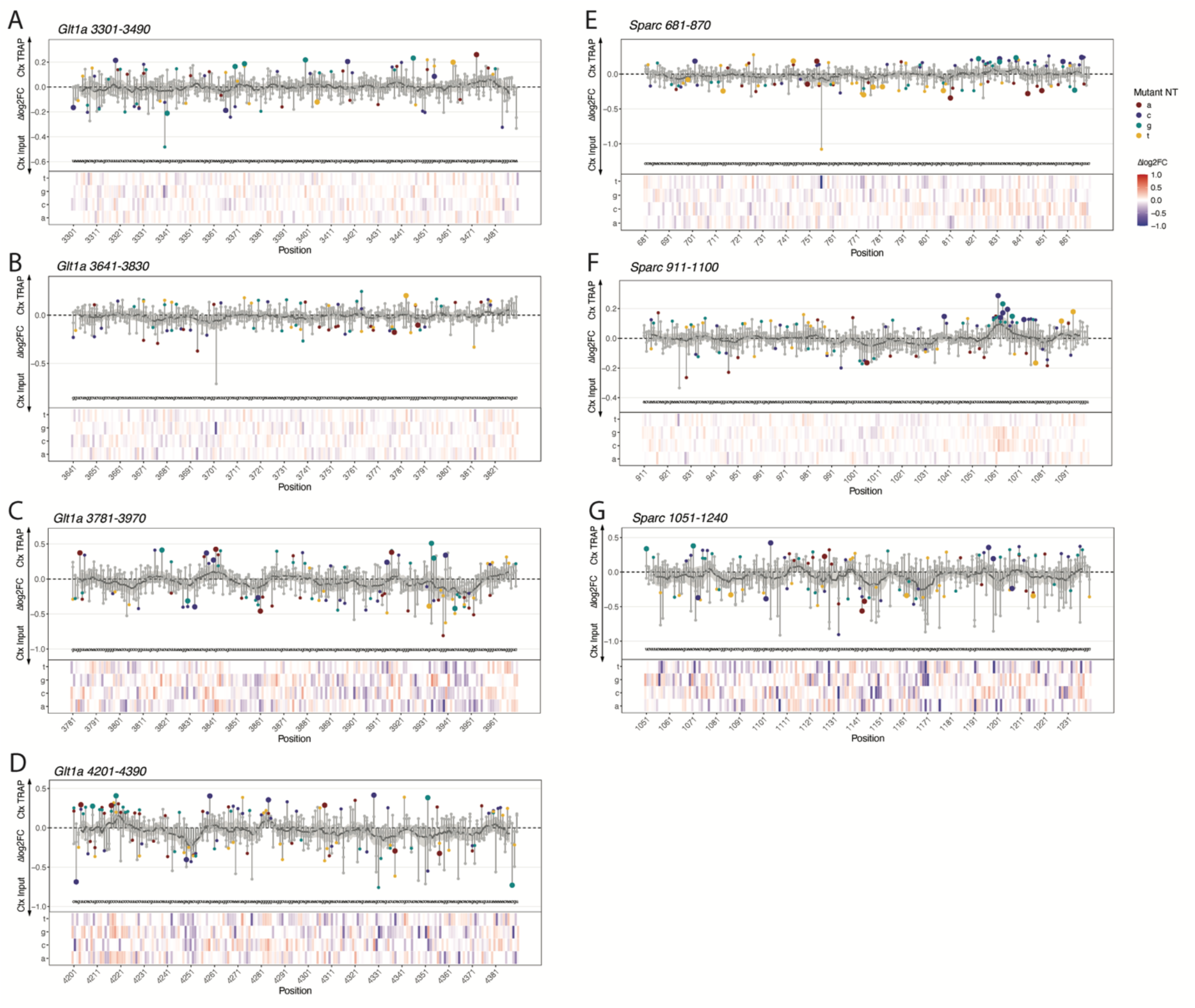
Single Nucleotide Mutagenesis Reveals Changes in Ribosome Occupancy. **A-G**. ΔRibosome Occupancy (log2FC(normCortex TRAP/normCortex input)(*y)* of mutagenesis elements (mutant nucleotide specified by red = A, purple = C, cyan = G, yellow = T/U) plotted against position of individual mutation in the 3’ UTR fragment *(x)* with heat map representation of the Δlog2FC below for **A**. *Glt1a*, positions 3301-3490, **B**. *Glt1a*, positions 3641-3830, **C**. *Glt1a*, positions 3781-3970, **D**. *Glt1a*, positions 4201-4390, **E**. *Sparc*, positions 681-870, and **F**. *Sparc*, positions 911-1100, and **G**. *Sparc*, positions 1051-1240. Significance determined after linear mixed model to account for random effect of replicate (Expression ∼ Fraction + (1|Replicate)) with FDR correction. Grey dots: ns; small colored dots: p < 0.05; larger colored dots: FDR < 0.05.

**Supplemental Figure S6:**
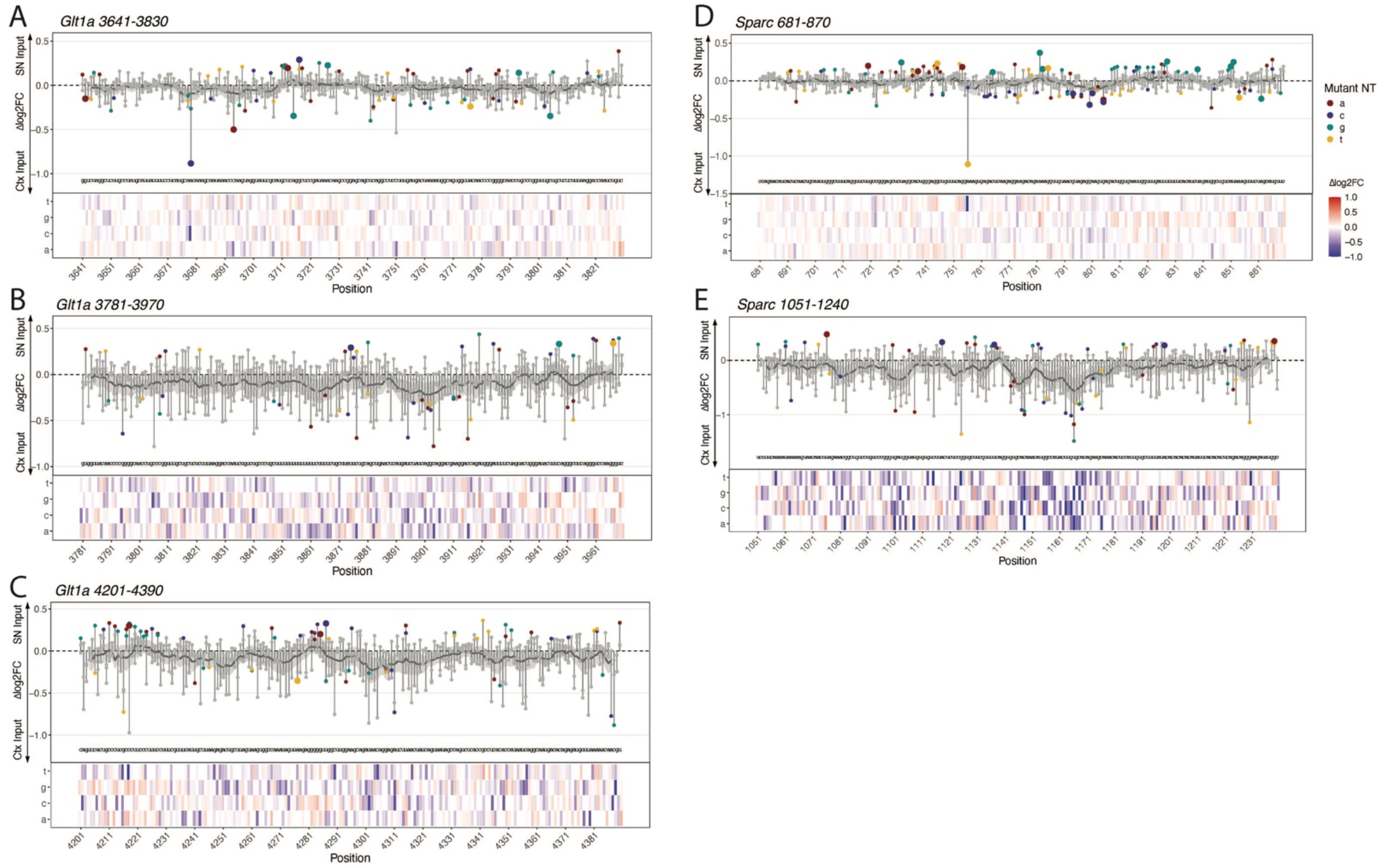
Single Nucleotide Mutagenesis Reveals Changes in Localization. **A-G**. ΔLocalization (log2FC(normSN input/normCortex input)(*y)* of mutagenesis elements (mutant nucleotide specified by red = A, purple = C, cyan = G, yellow = T/U) plotted against position of individual mutation in the 3’ UTR fragment *(x)* with heat map representation of the Δlog2FC below for **A**. *Glt1a*, positions 3641-3830, **B**. *Glt1a*, positions 3781-3970, **C**. *Glt1a*, positions 4201-4390, **D**. *Sparc*, positions 681-870, and **E**. *Sparc*, positions 1051-1240. Significance determined after linear mixed model to account for random effect of replicate (Expression ∼ Fraction + (1|Replicate)) with FDR correction. Grey dots: ns; small colored dots: p < 0.05; larger colored dots: FDR < 0.05.

**Supplemental Figure S7:**
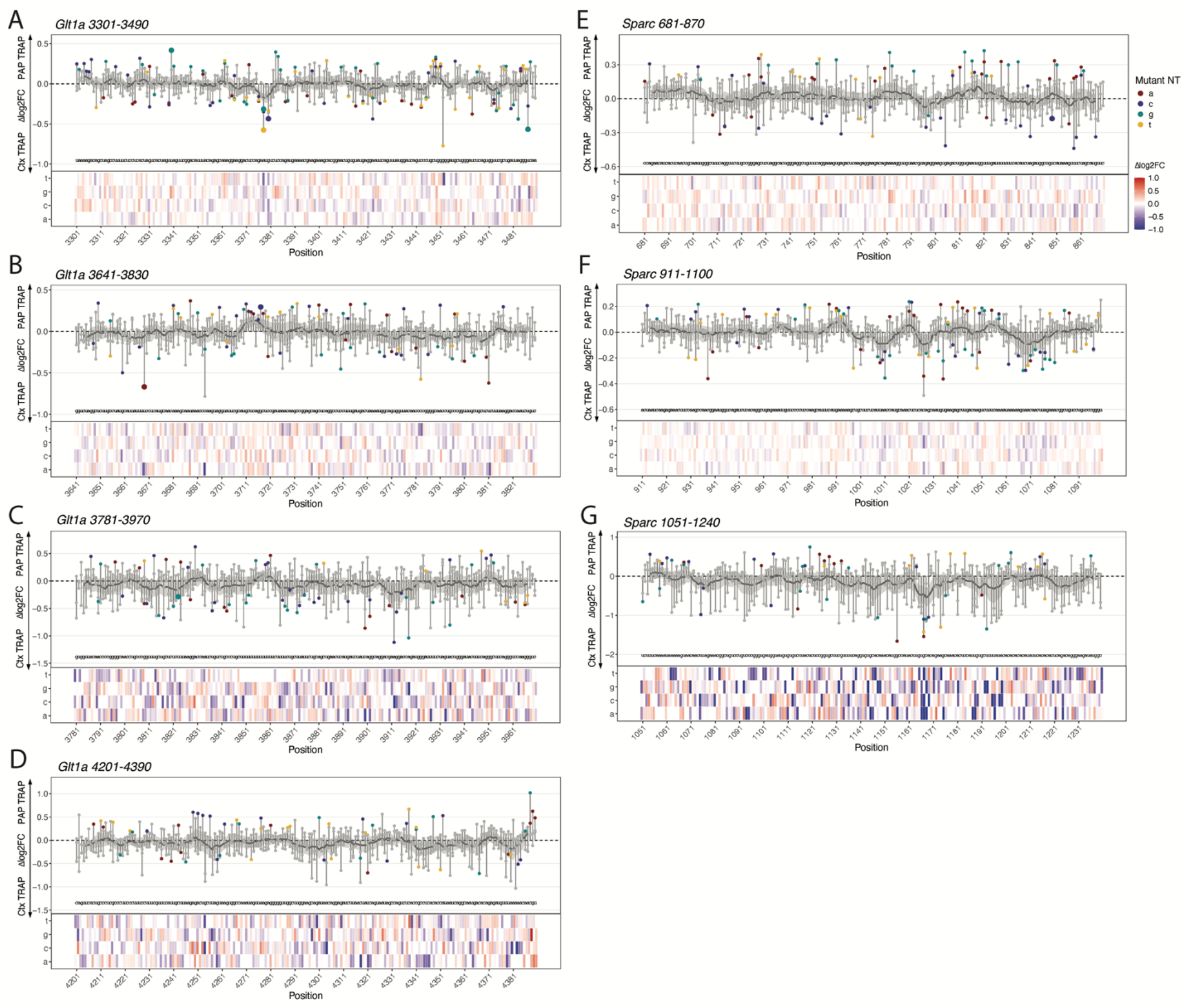
Single Nucleotide Mutagenesis Reveals Changes in Local Translation. **A-G**. ΔLocal Translation log2FC(normPAP-TRAP/normCortex TRAP)(*y)* of mutagenesis elements (mutant nucleotide specified by red = A, purple = C, cyan = G, yellow = T/U) plotted against position of individual mutation in the 3’ UTR fragment *(x)* with heat map representation of the Δlog2FC below for **A**. *Glt1a*, positions 3301-3490, **B**. *Glt1a*, positions 3641-3830, **C**. *Glt1a*, positions 3781-3970, **D**. *Glt1a*, positions 4201-4390, **E**. *Sparc*, positions 681-870, and **F**. *Sparc*, positions 911-1100, and **G**. *Sparc*, positions 1051-1240. Significance determined after linear mixed model to account for random effect of replicate (Expression ∼ Fraction + (1|Replicate)) with FDR correction. Grey dots: ns; small colored dots: p < 0.05; larger colored dots: FDR < 0.05.

**Supplemental Figure S8:**
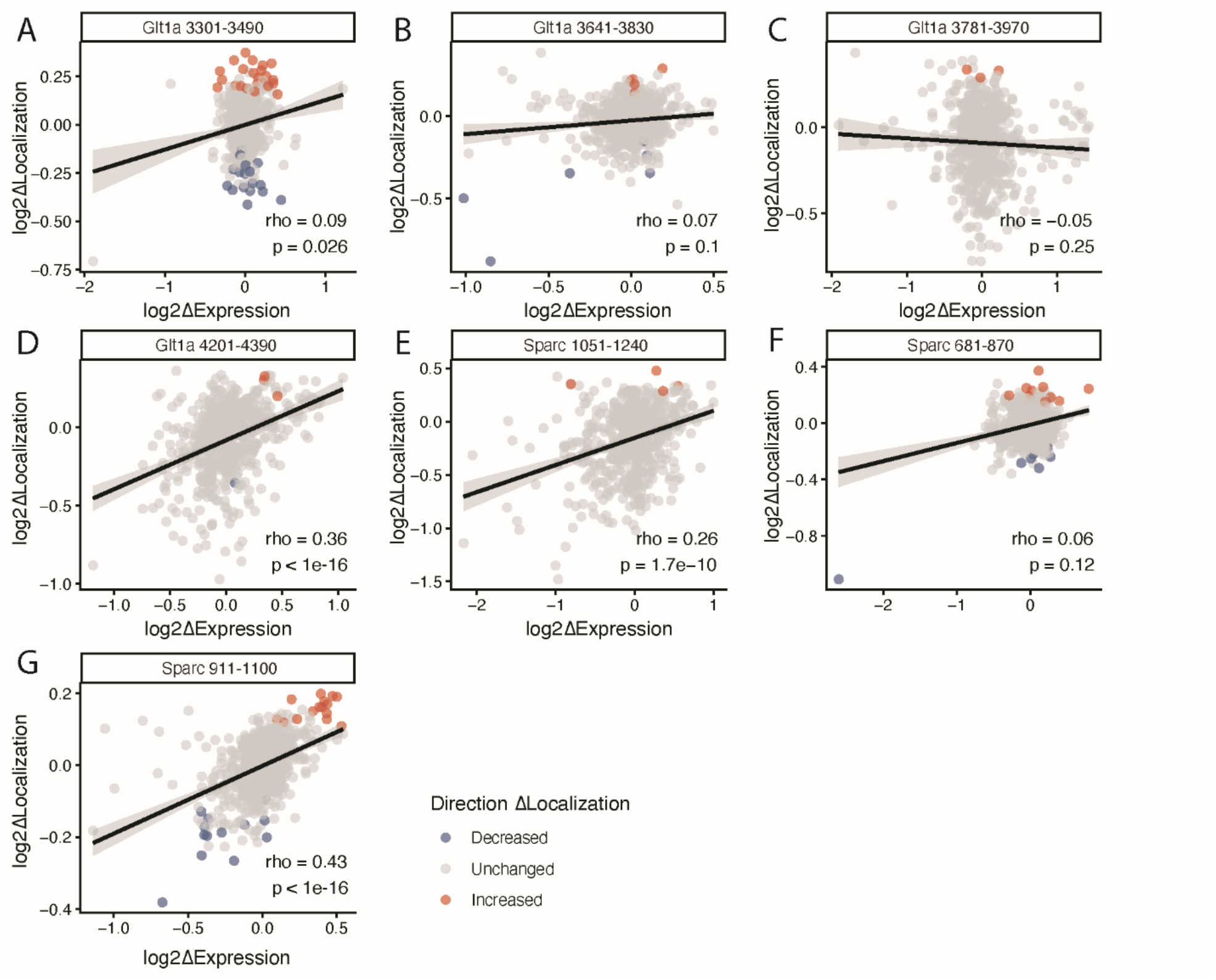
Correlations of ΔLocalization with ΔExpression. **A-G**. Spearman correlations of ΔExpression (log2(normCortex input/normAAV DNA)) and ΔLocalization (log2(normSN input/normCortex input)) with BH FDR adjusted p-value for **A**. *Glt1a* 3301-3490 (rho = 0.09; p = 0.026), **B**. *Glt1a* 3641-3830 (rho = 0.07; p = 0.1), **C**. *Glt1a* 3781-3970 (rho = -0.05; p = 0. 25), **D**. *Glt1a* 4201-4390 (rho = 0.36; p < 1e-16), **E**. *Sparc* 1051-1240 (rho = 0.26; p = 1.7e-10), **F**. *Sparc* 681-870 (rho = 0.06; p = 0.12), and **G**. *Sparc* 911-1100 (rho = 0.43; p < 1e-16). Dots colored based on significantly enriched (red), depleted (blue), or unchanged (gray) for measures of localization. Elements with FDR adjusted p < 1e-8 were considered significant.

**Supplemental Figure S9:**
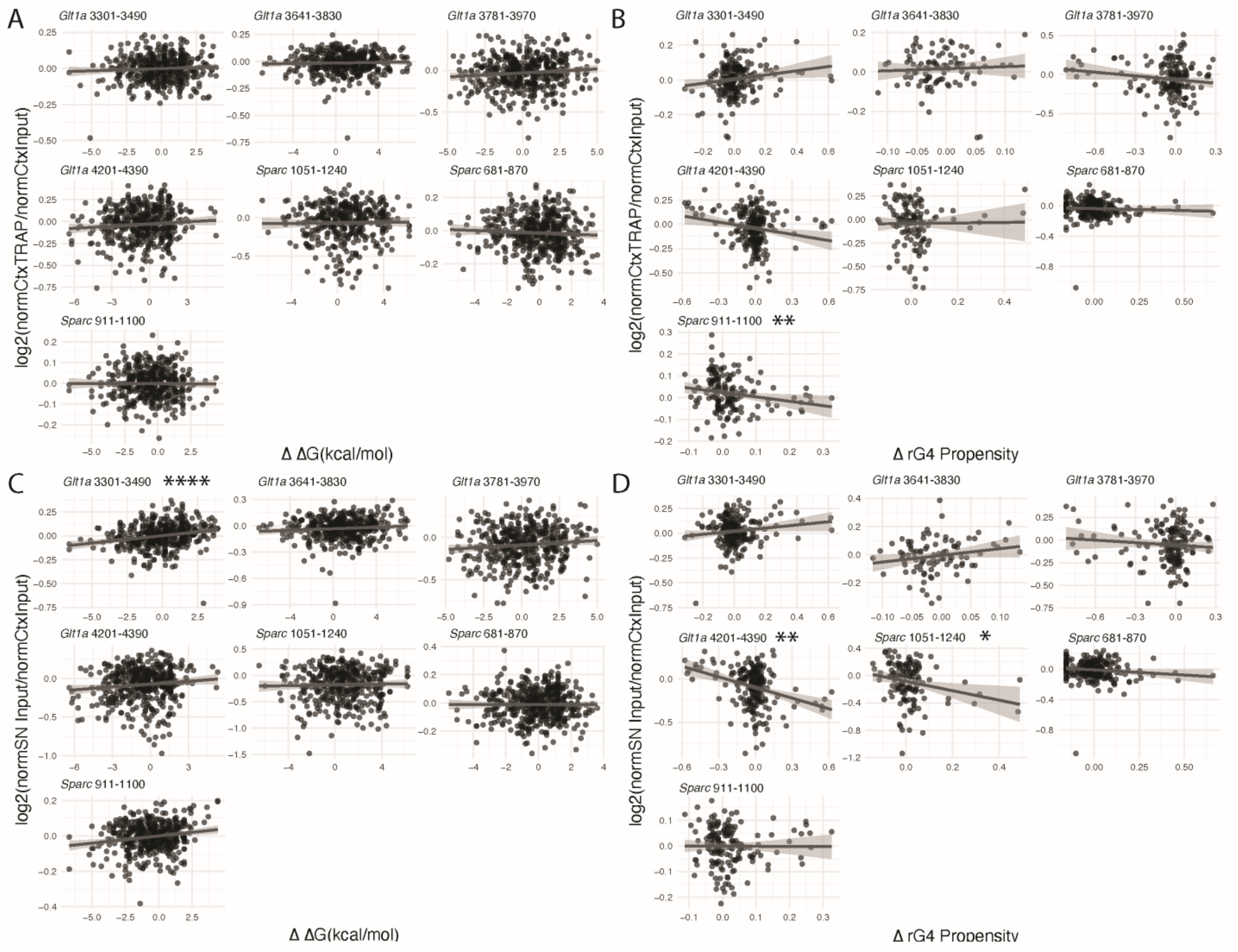
Correlations of ΔRibosome Occupancy and ΔLocalization with Changes in Mutagenesis Elements that Change RNA Secondary Structure and G-Quadruplex Propensity. **A-D**. For elements with mutations changing ΔG and rG4, **A-B**. Spearman correlations of each individual mutagenesis element’s **A**. ΔΔG or **B**. ΔrG4 with its ΔRibosome Occupancy (log2(normCortex TRAP/normCortex input)) separated by element group. **C-D**. Spearman correlations of each individual mutagenesis element’s **C**. ΔΔG or **D**. ΔrG4 with its ΔLocalization (log2(normSN Input/normCortex input)) separated by element group. Rho and FDR-corrected p values reported in Table 9. *FDR < 0.05, **FDR < 0.01, ****FDR < 1e-4

## Supplemental Tables

**Supplemental Table 1: SN-MPRA Library *Cis*-Regulatory Element Sequences**

A list containing each elements name, sequence, gene symbol, isoform number, gene ID, Chromosome, Position, Strand, start and end position, GC content, secondary structure prediction and DeltaG(kcal/mol).

**Supplemental Table 2: SN-MPRA Library Quality Control Information**

QC information for each fraction/replicate: number of counts, number of reads aligned to library, percent alignment, number of barcodes recovered, and percent of barcodes recovered.

**Supplemental Table 3: SN-MPRA Tiling Library Results**

Results summary for SN MPRA. Includes counts and CPMs for each fraction/replicate, logFC, p values, FDR values for each measure: ribosome occupancy (ctxtrap_ctxin), local ribosome occupancy (paptrap_snin), localization (snin_ctxin), and local translation (paptrap_ctxtrap).

**Supplemental Table 4: Dual Fluorescence Validation Constructs**

Sequences of validation constructs cloned into dual fluorescence reporters for post-natal electroporations and immunofluorescence.

**Supplemental Table 5: Primers for *Sparc* 3’UTR and *Sparc***Δ**661-890 cloning**

Primers used to generate Sparc 3’ UTR and *Sparc*Δ661-890 dual fluorescence validation constructs

**Supplemental Table 6: SN-MPRA Single Nucleotide Mutagenesis Library *Cis*-Regulatory Element Sequences**

A list containing each elements name, gene, sequence, 3’ UTR position start, 3’ UTR position end, original nucleotide, mutant nucleotide, sequence length, original GC content (of element group), element group, DeltaG (kcal/mol), and G-quadruplex (rG4) prediction score.

**Supplemental Table 7: Single Nucleotide Mutagenesis Library QC Metrics**

QC information for each fraction/replicate (sample): number of counts, number of reads aligned to library, percent alignment, number of elements recovered, and percent of elements recovered.

**Supplemental Table 8: SN-MPRA Single Nucleotide Mutagenesis Library Results**

Results summary for SN MPRA. Includes counts and CPMs for each fraction/replicate, logFC, p values, FDR values for each measure: normalized ribosome occupancy (ctxtrap_ctxin), normalized localization (snin_ctxin), and normalized local translation (paptrap_ctxtrap).

**Supplemental Table 9: Correlations between** Δ**Ribosome Occupancy**, Δ**Localization, and** Δ**Local Translation with Changes in RNA Secondary Structure, Conservation, and G-Quadruplex Propensity**

Pearson correlations and resulting FDR between ΔRibosome Occupancy, ΔLocalization, and ΔLocal Translation with change in RNA secondary structure (ΔΔG), phyloP score, and change in G-Quadruplex Propensity (ΔrG4).

**Supplemental Table 10: RBP Motifs that Overlap with Mutagenesis Regions Disrupting Localization**

List of RNA Binding Proteins with motifs that match regions of element groups in which mutagenesis significantly decreases localization

